# *α*_M_I-domain of Integrin Mac-1 Binds the Cytokine Pleiotrophin Using Multiple Mechanisms

**DOI:** 10.1101/2024.02.01.578455

**Authors:** Hoa Nguyen, Nataly P. Podolnikova, Tatiana P. Ugarova, Xu Wang

## Abstract

The integrin Mac-1 (α_M_β_2_, CD11b/CD18, CR3) is an important adhesion receptor expressed on macrophages and neutrophils. Mac-1 is also the most promiscuous member of the integrin family that binds a diverse set of ligands through its α_M_I-domain. However, the binding mechanism of most ligands is not clear. We have determined the interaction of α_M_I-domain with the cytokine pleiotrophin (PTN), a cationic protein known to bind α_M_I-domain and induce Mac-1-mediated cell adhesion and migration. Our data show that PTN’s N-terminal domain binds a unique site near the N- and C-termini of the α_M_I-domain using a metal-independent mechanism. However, stronger interaction is achieved when an acidic amino acid in a zwitterionic motif in PTN’s C-terminal domain chelates the divalent cation in the metal ion-dependent adhesion site of the active α_M_I-domain. These results indicate that α_M_I-domain can bind ligands using multiple mechanisms, and suggest that active α_M_I-domain prefers acidic amino acids in zwitterionic motifs.

**HIGHLIGHTS:**

- α_M_I-domain’s interaction with the cytokine pleiotrophin (PTN) was investigated with solution NMR.
- α_M_I-domain binds PTN using multiple mechanisms.
- PTN’s N-terminal domain binds both active and inactive α_M_I-domains using a unique site near α_M_I-domain’s termini.
- PTN’s C-terminal domain binds only active α_M_I-domain through a metal-dependent interaction.

## INTRODUCTION

Mac-1 (α_M_β_2_, CD11b/CD18, CR3) is a member of the αβ heterodimeric adhesion receptor family known as integrins. Mac-1 is primarily expressed in neutrophils, monocytes, and macrophages. It is responsible for many important activities in these cells, including phagocytosis, migration, and degranulation (Coxon et al., 1996; Lu and Springer, 1997; Ding et al., 1999; Prince et al., 2001). It has also been increasingly recognized as a vital regulator of the inflammatory state of these cells, capable of promoting both pro-inflammatory and anti-inflammatory pathways in a context dependent way (Rosetti and Mayadas, 2016). Mac-1’s diverse biological activities are closely connected with its broad ligand specificity. It is the most promiscuous member of the integrin family and is known to bind nearly 100 different ligands with little consensus among their structures (Lamers et al., 2021). The best-known and most characterized Mac-1 ligands include fibrinogen (Altieri et al., 1990; Ugarova et al., 1998), complement factor C3b (Michishita et al., 1993), and intercellular adhesion molecule 1 (ICAM-1) (Smith et al., 1989; Diamond et al., 1990), and GPIbα (Simon et al., 2000). Still, it is also known to bind such diverse ligands as JAM-3 (Santoso et al., 2002), elastase (Cai and Wright, 1996), myeloperoxidase (Johansson et al., 1997), plasminogen (Lishko et al., 2004), ovalbumin (Zhang and Plow, 1996), keyhole limpet hemocyanin (Shappell et al., 1990; Davis, 1992), CCN1(Schober et al., 2003), and CD40L(Wolf et al., 2011). Recently, it was also shown to bind SIRPα and mediate the merging of macrophages (Podolnikova et al., 2019). Although many ligands are known to bind Mac-1, the binding mechanisms of only a handful of ligands were confirmed with structures (Bajic et al., 2013; Jensen et al., 2016; Morgan et al., 2019; Trstenjak et al., 2020; Fernandez et al., 2022; Goldsmith et al., 2022). The binding interactions of other ligands and their functional consequences remain to be elucidated.

Surprisingly, despite the structural diversity among Mac-1 ligands, most bind to the I-domain of the α_M_ subunit (α_M_I-domain). α_M_I-domain is a two hundred residue Rossmann fold domain inserted between blades two and three of α_M_’s β-propeller. It contains a divalent metal binding site called the metal ion-dependent adhesion site (MIDAS). MIDAS facilitates metal ion-mediated binding of some ligands by allowing the divalent cation in the MIDAS to be chelated by an acidic amino acid in the ligand. The conformation of the α_M_I-domain is crucial to the strength of metal-mediated ligand binding. In particular, when the α_M_I-domain is in the inactive or “closed” conformation, the coordination of the metal in MIDAS does not allow efficient chelation of the metal ion by the ligand. However, when the α_M_I-domain is in the active or “open” conformation, the movement of its C-terminal helix away from MIDAS allows the ligand’s acidic amino acid to chelate the metal ion with much higher affinity (Lee et al., 1995).

Although metal-mediated ligand binding is prevalent among integrins, it has long been known that Mac-1 uses other ligand binding mechanisms because some Mac-1-binding sequence motifs do not contain acidic amino acids (Yakubenko et al., 2001). In addition, a systematic peptide screening revealed that motifs containing basic amino acids flanked by hydrophobic amino acids also have a high affinity for α_M_I-domain (Podolnikova et al., 2015b). This has led to the discovery of several new integrin ligands, including the cathelicidin peptide LL-37 (Lishko et al., 2016; Zhang et al., 2016), the opioid peptide dynorphin A (Podolnikova et al., 2015a), the chemokine PF4 (Lishko et al., 2018), the cytokine pleiotrophin (PTN) (Shen et al., 2017), and many cationic ligands (Podolnikova et al., 2015b).

PTN is a cytokine consisting of two thrombospondin type-1 repeat domains and plays important roles in cell differentiation and proliferation (Ryan et al., 2016; Wang, 2020). It is also a modulator of microglia activity (Fernandez-Calle et al., 2017; Miao et al., 2019; Linnerbauer et al., 2021), where Mac-1 is highly expressed. Our previous work showed that PTN induced adhesion and migration of various Mac-1-expressing cells, and PTN can activate ERK1/2 in a Mac-1-dependent manner (Shen et al., 2017). As a representative cationic ligand, we wondered if PTN’s interaction with α_M_I-domain can provide insight into the binding mechanism of this class of ligands. Solution NMR characterization of the interaction of PTN with inactive α_M_I-domain showed that the interaction is independent of Mg^2+^ and the binding interface includes the α5-β5 loop of inactive α_M_I-domain and PTN’s N-terminal domain (PTN-NTD) (Feng et al., 2021). However, these data do not explain the observation that Mg^2+^ ions significantly enhanced the binding of PTN to active α_M_I-domain (Shen et al., 2017), implying that a divalent cation-dependent mechanism is also part of the interaction.

In the current study, we investigated PTN’s interaction with both active and inactive α_M_I-domain. Our data indicate the Mg^2+^-independent interactions between α_M_I-domain and PTN involve a binding interface formed by PTN-NTD and residues from the α5-β5/α6-β6 loops near the termini of α_M_I-domain. However, the Mg^2+^-dependent mechanism uses acidic amino acids in PTN to chelate the metal ion in α_M_I-domain’s MIDAS. In particular, residue E98 in the PTN’s C-terminal domain (PTN-CTD) was shown to be the preferred chelator of MIDAS metal. We attribute this preference to the fact that E98 is part of a zwitterionic motif and its interaction with the metal is stabilized by favorable electrostatic interactions between charged residues around E98 and the MIDAS. Using these data, we created models of the inactive α_M_I-domain-PTN-NTD and active α_M_I-domain-PTN-CTD complexes using HADDOCK (Dominguez et al., 2003). We also carried out molecular dynamics (MD) simulations of these models to provide additional insights into the mechanism by which the α_M_I-domain can interact with its ligands.

## RESULTS

### Mg^2+^-independent interactions between PTN and α_M_I-domain

We have shown previously that although PTN has the highest affinity for active α_M_I-domain, it also interacts weakly with inactive α_M_I-domain (Shen et al., 2017). Additional studies showed the metal-independent interaction is mediated by the α5-β5 loop near the termini of inactive α_M_I-domain and PTN-NTD (Feng et al., 2021). To determine the structure of the inactive α_M_I-domain-PTN-NTD complex, we collected the F1-^13^C-edited/F3-^13^C,^15^N-filtered HSQCNOESY spectrum of a sample containing 0.2 mM ^13^C, ^15^N-labeled inactive α_M_I-domain and 1.0 mM unlabeled PTN-NTD with no Mg^2+^. These data revealed many intermolecular contacts between PTN-NTD and inactive α_M_I-domain (Figure 1). In particular, methyl groups of residue L32 in PTN-NTD had definitive contacts with G263 and I265 in the α5-β5 loop of inactive α_M_I-domain. In addition, the methyl group of residue T26 in PTN-NTD contacted residue K290 in inactive α_M_I-domain, T34 in PTN-NTD contacted both residues K290 and P291 in inactive α_M_I-domain, and T50 in PTN-NTD contacted residue P291 in inactive α_M_I-domain. R52 in PTN-NTD also contacted I265 in inactive α_M_I-domain. To confirm the assignments of the PTN residues, we collected the F1-^13^C,^15^N-filtered/F3-^13^C-edited NOESYHSQC spectrum of a sample containing 0.2 mM unlabeled inactive α_M_I-domain and 0.5 mM ^13^C, ^15^N-labeled PTN-NTD. These data provided the ^13^C chemical shifts of the PTN-NTD atoms at the interaction interface (Figure S1). It should be noted that the NMR signal from K290 was mis-assigned to K168 previously (Feng et al., 2021). In the current study, the assignment of residue K290 in α_M_I-domain was confirmed through selective ^15^N-labeling of lysines and a K290R mutant of inactive α_M_I-domain (Figure S2). The ^15^N-edited NOESYHSQC spectrum of inactive α_M_ I-domain containing selectively ^15^N-labeled lysines allowed the side chain hydrogens of lysines to be assigned. K290 was the only lysine whose side chain proton resonance frequencies matched the intermolecular NOE cross peaks in the F1-^13^C-edited/ F3-^13^C,^15^N-filtered HSQCNOESY spectrum. We also acquired an F1-^13^C-edited/F3-^13^C,^15^N-filtered HSQCNOESY spectrum of ^13^C,^15^N-labeled inactive α_M_I-domain in the presence of PTN-CTD. The data produced no identifiable intermolecular cross peaks between PTN-CTD and inactive α_M_I-domain (Figure S3). This is consistent with the observation that only PTN-NTD can produce the large chemical shift perturbations in ^15^N-HSQC spectrum of inactive α_M_I-domain while PTN-CTD induces only minor perturbations. (Feng et al., 2021).

**Figure 1.**
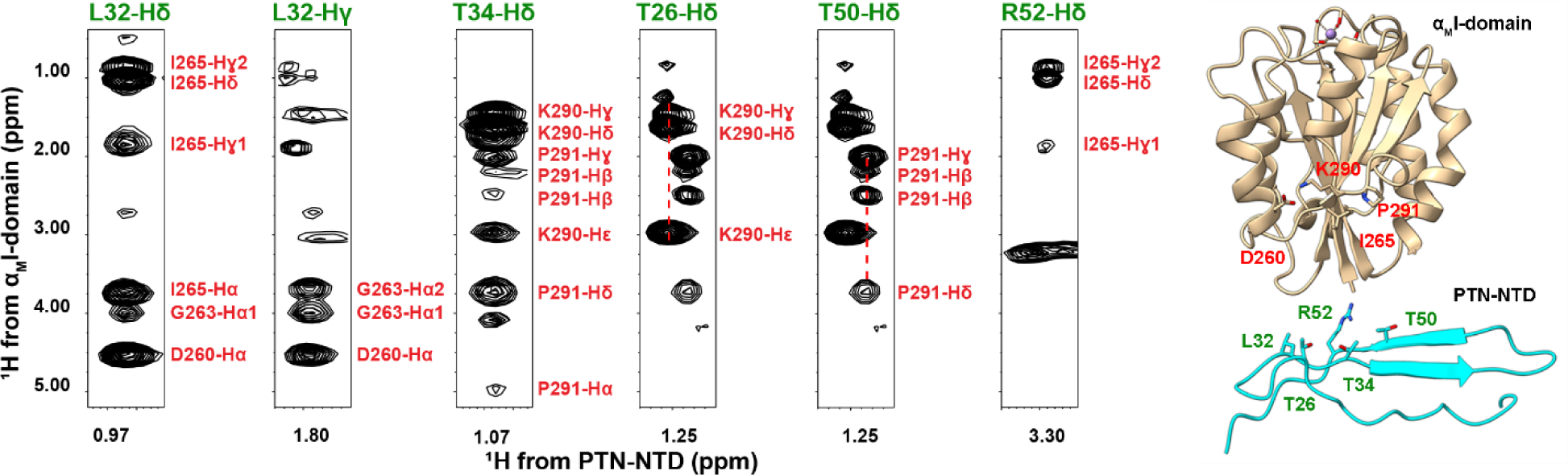
Contacts between the inactive α_M_I-domain and PTN-NTD. Strips from F1-^13^C-edited/F3-^13^C,^15^N-filtered HSQCNOESY spectrum of ^13^C,^15^N-labeled inactive α_M_I-domain and unlabeled PTN-NTD. Inactive α_M_I-domain assignments are shown in red. PTN-NTD assignments are shown in green. Ribbon diagram of inactive α_M_I-domain and PTN-NTD with residues involved in the intermolecular contacts labeled are shown on the right.

Altogether, these data indicate that the interaction interface between PTN-NTD and inactive α_M_I-domain includes the α5-β5 and α6-β6 loops of inactive α_M_I-domain and PTN-NTD. In addition, both inactive and the Q163C/Q309C active α_M_I-domain (Shimaoka et al., 2002) induced similar chemical shift changes in PTN-NTD (Figure S4). This implies that PTN-NTD most likely has similar metal-independent interactions with active α_M_I-domain, suggesting that the Mg^2+^-independent interaction between PTN and α_M_I-domain is not sensitive to the activation state of α_M_I-domain.

### Mg^2+^-dependent interactions between α_M_I-domain and PTN

Our previous study has shown that PTN’s affinity for active α_M_I-domain is higher than for inactive α_M_I-domain and Mg^2+^ is required for the interaction (Shen et al., 2017). To elucidate the underlying mechanism, we investigated PTN’s interaction with the Q163C/Q309C mutant of α_M_I-domain, which is forced into the active conformation by a well-placed disulfide bond (Shimaoka et al., 2002). A previous study has shown that the Q163C/Q309C mutant is more suitable for NMR studies because, while it has the same conformation and ligand affinity as active α_M_I-domains created by the removal of residue I316 (Xiong et al., 2000), it possesses higher stability and better NMR spectral quality (Nguyen et al., 2023). Because of these favorable properties, the Q163C/Q309C mutant was chosen for this study. All subsequent mentions of active α_M_I-domain in this article refer to the Q163C/Q309C mutant.

Interestingly, when we titrated the active α_M_I-domain with PTN in the presence of Mg^2+^, PTN induced similar spectral changes in active α_M_I-domain as glutamate, a ligand known to chelate the metal in the MIDAS of the I316G active α_M_I-domain (Vorup-Jensen et al., 2005) (Figure 2A). In particular, adding either glutamate or PTN to the Mg^2+^-bound active α_M_I-domain led to intensity decreases in some signals and the appearance of new signals, consistent with slow time scale exchange between ligand-free and ligand-bound forms of α_M_I-domain. Figure 2B shows the signal of residue G228 of α_M_I-domain undergoing slow exchange when titrated with PTN. Three of the new signals that appeared in the presence of ligands were assigned to residues G143, S144, and I145, all are residues in one of the MIDAS segments that directly chelate the metal. The similarity in ligand-induced spectral perturbations shows that glutamate and PTN interacted with the MIDAS similarly. This finding supports the idea that PTN binds active α_M_I-domain through metal-mediated interactions. Some MIDAS residues also exhibit PTN-domain-specific chemical shifts. In particular, wild-type PTN binding produced two signals from residue S144. However, only one of the signals was seen when PTN-CTD was the ligand whereas PTN-NTD only produced the other signal (Figure S5). We interpret this as an indication that chelation of the metal by different domains resulted in differences in the chemical shifts of the S144 signal. The fact that both signals are present when wild-type PTN is the ligand indicates both PTN-NTD and PTN-CTD can chelate the metal. However, PTN-CTD produced higher intensity signals that are consistent with stable chelation of the MIDAS metal. In contrast, the intensities of these signals were much weaker when PTN-NTD was mixed with active α_M_I-domain (Figure 2A). This indicates PTN-CTD may have higher affinity for active α_M_I-domain than PTN-NTD. It should also be noted that PTN did not induce these changes in the absence of Mg^2+^ (Figure S6), supporting the idea that the observed active α_M_I-domain spectral changes induced by PTN are metal-dependent.

**Figure 2.**
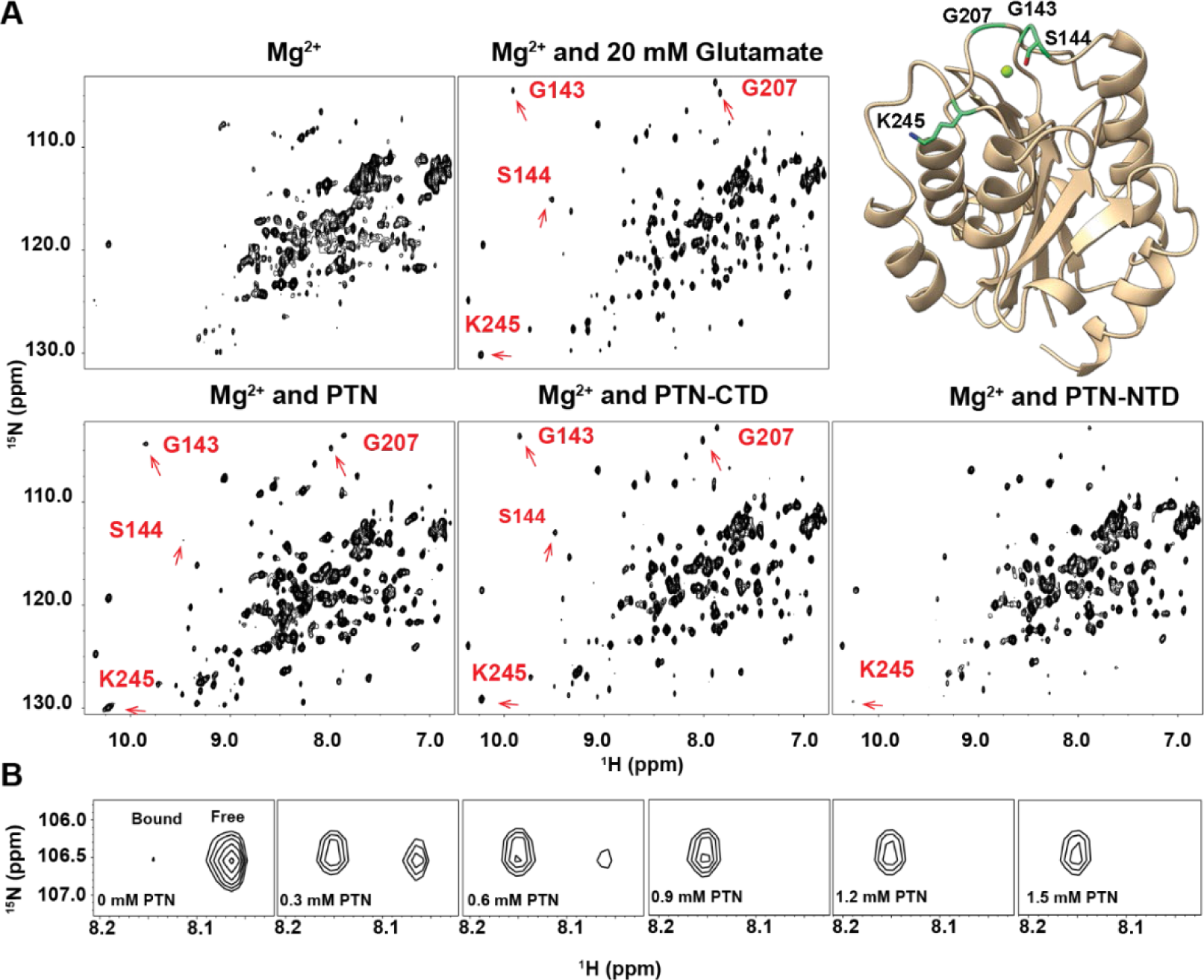
Ligand-induced changes in the ^15^N-HSQC spectrum of active α_M_I-domain. (A) ^15^N-HSQC of the active α_M_I-domain in the presence of different ligands. The changes in the spectrum of active α_M_-I domain produced by wild-type PTN or PTN domains are similar to those produced by glutamate, including the appearance of the residues around MIDAS (G143, S144, G207, K245). The signal intensities of the MIDAS residues induced by PTN-NTD were much weaker than those induced by PTN-CTD at the same concentration. Ribbon representation of active α_M_I-domain with labeled MIDAS residues are shown on the top right. B) ^15^N-HSQC signal of residue G228 in Mg^2+^ -saturated active α_M_I-domain with different concentrations of PTN. The signal underwent slow time scale exchanges when titrated with PTN.

Using intensity increases in the ligand-bound species as a measure of the binding allowed us to obtain the *K*_d_ of interaction f by fitting the intensity changes to a one-to-one binding model. We also carried out principle component analysis on the spectral data using the software TRENDNMR (Xu and Van Doren, 2016) and used the magnitude of principle component 1 as a measure of the binding to estimate the *K*_d_ of interaction. Figure 3A and Table S1 show *K*_d_ s obtained by these analyses. The *K*_d_ for glutamate was ∼ 5.5 mM, whereas the *K*_d_ of interaction for PTN was ∼ 0.1 mM, significantly lower than that of the metal-independent interaction (∼ 1 mM) (Feng et al., 2021). To determine which domain of PTN is responsible for the metal-dependent interaction, we titrated active α_M_I-domain with PTN-CTD and PTN-NTD. The *K*_d_ for PTN-CTD binding was ∼ 0.1 mM. The *K*_d_ for PTN-NTD binding could not be estimated accurately because of the low intensities of the new ligand-induced signals, but it is at least 9 times greater than the *K*_d_ for PTN and PTN-CTD. These results support the conclusion that PTN-CTD contributed more to binding active α_M_I-domain through metal-chelation.

**Figure 3:**
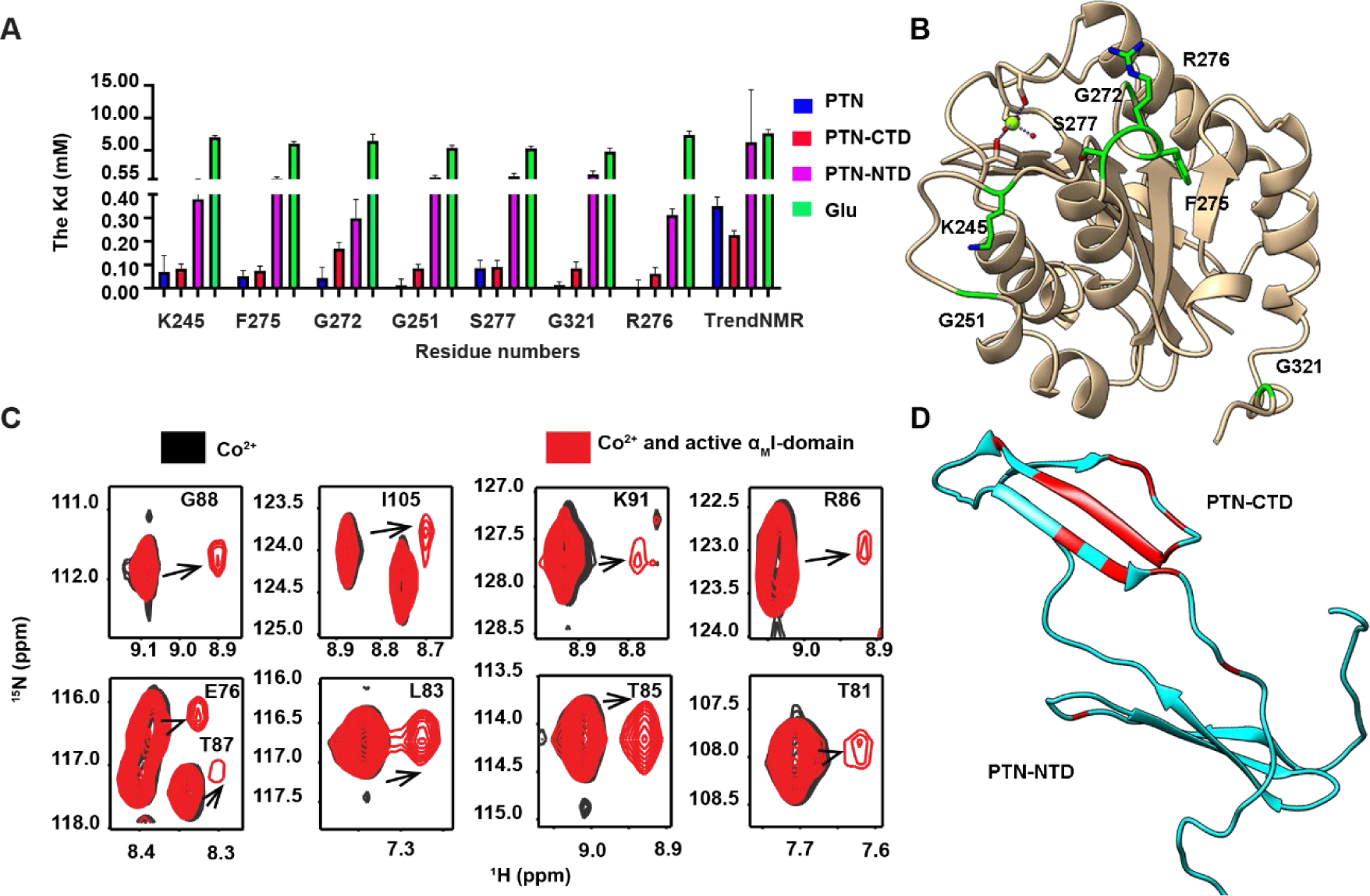
PTN-CTD is the binding site for active α_M_I-domain. A) Active α_M_I-domain’s *K*_d_ of binding for wild-type PTN (blue), PTN-CTD (red), PTN-NTD (magenta), and glutamate (green). *K*_d_s were calculated by either fitting the ligand induced signal intensity increases in six residues with high signal-to-noise or using global spectral changes estimated with TrendNMR. Error bars reflect S.D. in data fitting. B) The ribbon representation of active α_M_I-domain with residues used to calculate the *K*_d_ shown in the stick form. C) ^15^N-HSQC of PTN in the presence of Co^2+^ (black) and Co^2+^/active α_M_I-domain (red). D) The ribbon representation of PTN with residues exhibiting PCS shown in red. All except two residues exhibiting a PCS peak were located in PTN-CTD.

Another observation supporting PTN-CTD as the dominant metal binding domain is that Co^2+^-bound active α_M_I-domain induced pseudocontact shifts (PCS) mostly in residues from PTN-CTD. The MIDAS of α_M_I-domain can chelate paramagnetic Co^2+^ and Co^2+^ bound α_M_I-domain retains its ligand affinity (Michishita et al., 1993). The dipole-dipole interaction between the paramagnetic Co^2+^ ion and nearby atoms can induce a change in the chemical shifts of these atoms, commonly referred to as PCS (Nitsche and Otting, 2017). In protein-ligand interactions, the binding of the ligand to a paramagneti metal-containing protein can induce PCS in the ligand, if the interaction is sufficiently rigid. To investigate whether these transferred PCS can be observed in PTN, we collected the spectrum of ^15^N-labeled PTN in the presence and absence of Co^2+^-bound active α_M_I-domain. The results showed that the presence of Co^2+^-bound active α_M_I-domain induced an additional signal from some PTN residues (Figure 3C). The chemical shift differences between the new and original signals were consistent with diagonal shifts expected of small PCS. Most residues exhibiting a PCS peak were in PTN-CTD (Figure 3D). To confirm these signals resulted from PTN-CTD’s binding to Co^2+^-bound active α_M_I-domain, we also collected similar data of PTN-CTD with Mg^2+^-bound active α_M_I-domain and Co^2+^-bound inactive α_M_I-domain. The absence of either Co^2+^ or active α_M_I-domain produced no PCS in PTN-CTD (Figure S7). These results support the conclusion that PTN-CTD maintains more stable interactions with active α_M_I-domain. It should be noted that the intensities of these PCS signals are only about 11 % of the non-PCS signals, and higher concentrations of α_M_I-domain or Co^2+^ did not increase the relative intensities of the PCS signals (data not shown). This implies that PTN-CTD may bind to active α_M_I-domain in multiple ways and only some binding modes can induce PCS.

To identify which residue in PTN-CTD chelates the divalent cation in MIDAS, we collected the F1-^13^C-edited/F3-^13^C,^15^N-filtered HSQCNOESY spectrum of ^13^C-labeled active α_M_I-domain and unlabeled PTN-CTD. However, no intermolecular NOE was observed (Figure S8). This indicates no significant contact exists between the side chains of these proteins. We then collected ^1^H-^1^H and ^1^H-^13^C projections of the ^13^C-HSQC-NOESY-^15^N-HMQC spectrum of a sample containing 1 mM ^13^C-labeled PTN and 0.25 mM ^2^H, ^15^N-labeled active α_M_I-domain. Our data showed that several MIDAS residues from active α_M_I-domain, including G143, S144, I145, and R208, have intermolecular contacts with a glutamate side chain and the side chain methyl group of A93 in PTN-CTD (Figure 4). Due to chemical shift degeneracy in glutamate side chain atoms, we could not identify the exact glutamate. However, the closest glutamate to A93 is E98. Therefore, we hypothesized that E98 was likely the chelator of the metal ion in MIDAS. We also collected 4D ^13^C-HSQC-NOESY-^15^N-HMQC spectrum of ^2^H, ^15^N-labeled PTN-CTD in the presence of ^13^C-labeled active α_M_I-domain. However, no intermolecular contacts were detected. These results indicate that while backbone amide hydrogens of active α_M_I-domain were at the binding interface, none of the backbone amide hydrogens in PTN-CTD were at the interface.

**Figure 4.**
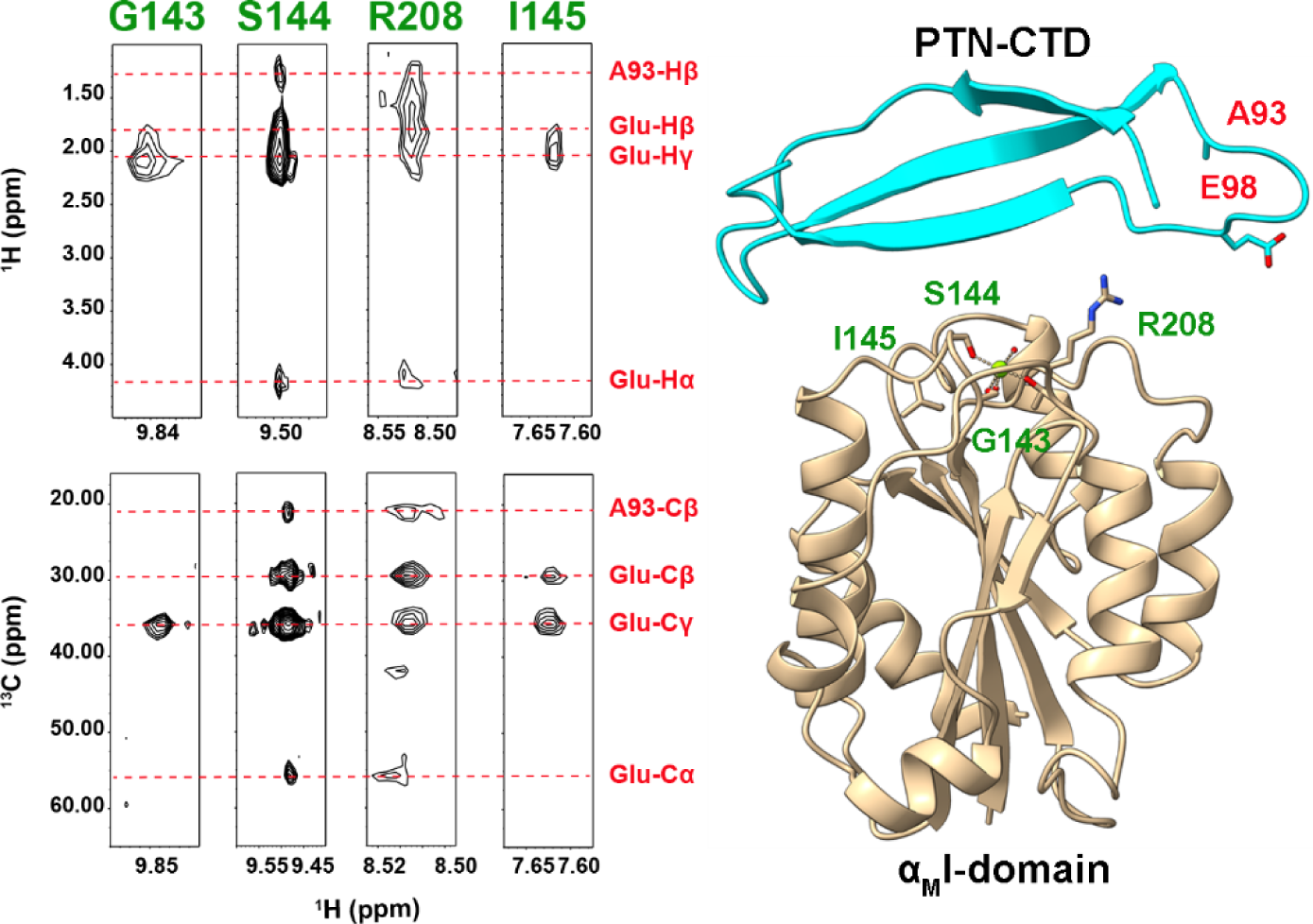
Contacts between backbone amide hydrogen of ^2^H, ^15^N-labeled active α_M_I-domain and ^13^C-labeled PTN-CTD seen in ^1^H-^1^H and ^1^H-^13^C projections of 4D ^13^C-HSQC-NOESY-^15^N-HMQC. The assignments of PTN-CTD atoms are labeled in red and the assignments of α_M_I-domain atoms are labeled in green. Ribbon representations of active α_M_I-domain and PTN-CTD with residues involved in the intermolecular contacts labeled are shown on the right. Due to degenerate chemical shifts, the glutamate cannot be assigned unambiguously.

To confirm that E98 is involved in metal chelation, we prepared several mutants of PTN-CTD. There are three acidic clusters in PTN-CTD, including E76/D78, E66/E98, and a string of four acidic amino acids in the unstructured C-terminal tail (E120/E127/E132/D136) (Figure 5B). We created PTN-CTD mutants missing one of the three clusters. We also mutated H95, which is in the 90s loop (residues R92 to K101 in PTN) and can potentially chelate metal ions. To monitor the binding, we titrated active α_M_I-domain with different PTN-CTD mutants and estimated the *K*_d_ using signal intensity increases experienced by the ligand-bound species. The results show that the mutation of E98 reduced the affinity most significantly (Figure 5A and Table S2). In particular, the removal of E76/D78 and the C-terminal tail (PTN-CTD Δtail) had only a marginal effect on PTN-CTD affinity whereas the mutation of E98 alone led to more than 5 fold decrease in affinity. Although the H95S mutation did not change the affinity drastically, both H95S and E98Q mutations produced large changes in chemical shift perturbation patterns in active α_M_I-domain when compared to wild-type PTN-CTD (Figure 5B and Table S3). This indicates that E98 and H95 are in the binding interface. It is worth noting that the mutation of E98 alone was not sufficient to eliminate the binding. The mutation of other acidic clusters also produced small decreases in affinity, indicating that other acidic amino acids also act as the chelator.

**Figure 5.**
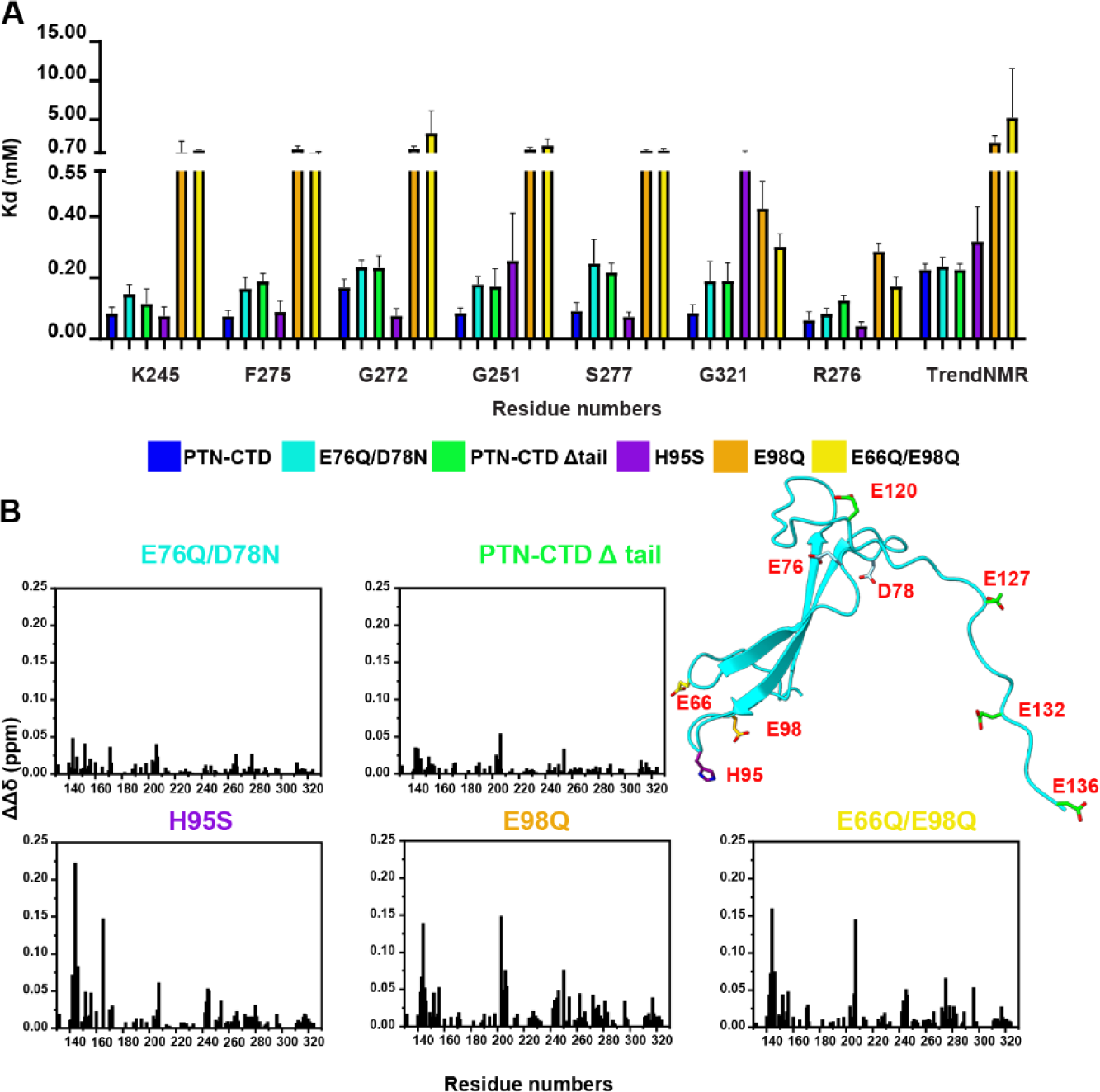
The effect of PTN mutations on its interactions with active α_M_I-domain. (A) The *K*_d_ of the interaction between active α_M_I-domain and PTN-CTD mutants. The *K*_d_s were obtained by fitting the ligand induced intensity changes of either signals with high signal-to-noise ratio or by fitting the global spectral changes estimated using TRENDNMR. *K*_d_ for wild-type PTN-CTD is shown in blue, E76Q/D78N is shown in cyan, PTN-CTD Δtail is shown in lime, H95S is shown purple, E98Q is shown in orange, E66Q/E98Q is shown in yellow. Error bars reflect S. D. in data fitting. (B) Differences in active α_M_I-domain backbone amide chemical shift changes induced by PTN-CTD mutants (ΔΔδ). The values were calculated by subtracting the chemical shift changes induced by the PTN-CTD mutant from the chemical shift changes induced by wild-type PTN-CTD. The ribbon representation of PTN-CTD with the mutated residues in the stick representation is shown on the right.

In addition to observing the effects of PTN-CTD mutations on the ^15^N-edited HSQC spectrum of active α_M_I-domain, we also examined the effect of H95 and E98 mutations on the PCS induced in PTN-CTD by Co^2+^-bound active α_M_I-domain. Figure 6A shows the impact of the mutations on the PCS signals of PTN-CTD residues with the strongest PCS peaks. The data revealed that the E98Q and H95S mutations diminished the PCS signal intensities of these residues by more than 75 %. The effect of the mutation of another acidic cluster, E76Q/D78N, on the PCS was far smaller. In particular, the PCS signal of W52 side chain indole Nε-Hε was not changed at all by the E76Q/D78N mutations. These data imply that both E98 and H95 are important to maintaining stable interactions needed to produce PCS signals.

**Figure 6.**
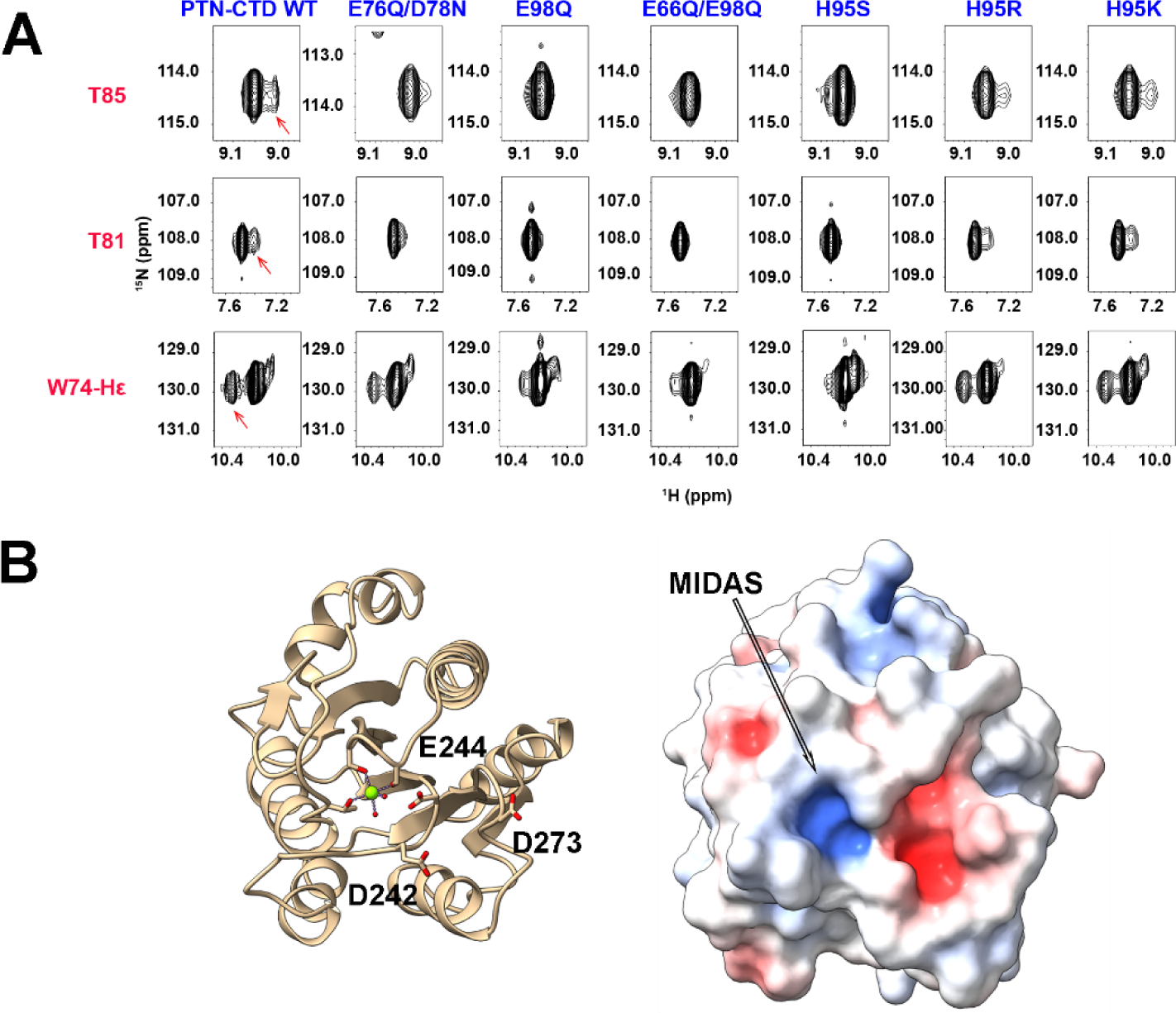
The role of H95 in binding active α_M_I-domain. A) Effect of PTN-CTD mutations on PCS induced by Co^2+^-bound active α_M_I-domain. The mutation of E98Q, E66Q/E98Q and H95S drastically reduced the PCS of PTN-CTD residues. PCS signals of residues are indicated by red arrows. B) Ribbon (left) and surface (right) representations of active α_M_I-domain. The Mg^2+^ ion in MIDAS is represented by the green sphere. Side chains of amino acids in the acidic patch are shown and labeled. Surface is colored based on the electrostatic surface potential range of -10 *k*_B_T/e (red) to 10 *k*_B_T/e (blue). Two representations are shown in the same orientation.

One question is how H95 interacts with MIDAS. We hypothesized that H95 in PTN-CTD interacts with the acidic pocket formed by α_M_I-domain residues D242, E244, and D273 next to the MIDAS (Figure 6B), thereby stabilizing the interactions between E98 and active α_M_I-domain. Similar stabilizing interactions between ligands and MIDAS of α I-domains were seen in the interactions of leukocidin GH and GP1bα with α_M_I-domain as well as in the binding of ICAM-1 to α_L_I-domain (Shimaoka et al., 2003; Morgan et al., 2019; Trstenjak et al., 2020). To confirm this, we prepared H95K and H95R mutants of PTN-CTD. Replacing H95 with another basic amino acid would preserve the electrostatic interaction with the acidic pocket near MIDAS and the PCS resulting from this more stable interaction would be retained. Figure 6A shows the ^15^N-HSQC spectra of H95K and H95R mutants of PTN-CTD in the presence of Co^2+^-bound active α_M_I-domain. The data demonstrate that, unlike the H95S mutation, substituting H95 with a basic amino acid preserves the PCS.

### Modeling of the complexes formed by α_M_I-domain and PTN domains

To model both Mg^2+^-dependent and Mg^2+^-independent interactions between PTN and α_M_I-domain, we docked PTN domain structures onto the structure of either active or inactive α_M_I-domain. We first confirmed that the structure of α_M_I-domain was not changed significantly by PTN domains. To do this, we assessed the conformation of the PTN domain-bound α_M_I-domain using the PCS induced in α_M_I-domain by Co^2+^. Our data show that the PCS of ligand-bound forms of both inactive and active α_M_I-domain fit the crystal structures of free α_M_I-domain well. In particular, the 81 PCS measured for the PTN-NTD bound inactive α_M_I-domain fitted the crystal structure of inactive α_M_I-domain (PDB ID 1JLM) with a Q factor of 0.055, and the ∼ 30 PCS measured for the PTN-CTD bound active α_M_I-domain fitted the crystal structure of active α_M_I-domain (PDB ID 1IDO) with a Q factor of 0.071 (Figure S9). These data indicate that the crystal structures of both inactive and active α_M_I-domain were a good starting point for modeling. For inactive α_M_I-domain, we also obtained the Cα chemical shifts in the presence and absence of PTN-NTD. These chemical shifts are excellent predictors of secondary structures of the protein (Wishart et al., 1991, 1992; Wishart and Sykes, 1994). Analysis of the Cα chemical shifts showed the secondary structure of the protein has not changed (Figure S10). We also collected the backbone amide ^1^H-^15^N residual dipolar couplings (RDC) of α_M_I-domain aligned in neutral polyacrylamide gel, both in the presence and absence of PTN-NTD. Although PTN-NTD appears to change the alignment of α_M_I-domain, both sets of RDCs fit the crystal structure of inactive α_M_I-domain (PDB code 1JLM) well (Conilescu Q factor of 0.25 and 0.28, respectively) (Figure S11). Because each PTN domain is small and stabilized by multiple disulfide bonds, we do not expect interactions with α_M_I-domain to change their structures.

To construct the model, we docked PTN-NTD onto inactive α_M_I-domain using the program HADDOCK (Dominguez et al., 2003). The intermolecular NOEs were included as non-ambiguous distance constraints (Table S4). Loop residues identified as being in the interface (residues 260 to 266, 289 to 293 in α_M_I-domain, and residues 25 to 27, 32 to 34, and 46 to 52 in PTN-NTD) were designated as flexible in the docking. The crystal structure of inactive α_M_I-domain (PDB 1JLM) and the NMR structure of PTN-NTD (PDB 2N6F) were used as the starting structures. Clustering analysis of the 200 resulting models showed all models belonged to the same cluster. This indicates the NOE information is sufficient to determine the structure unambiguously. After superimposing the inactive α_M_I-domain, the backbone RMSD of the structured region of PTN-NTD (residues 16 to 56) among the top 10 structures with the lowest overall HADDOCK scores was 1.9 Å. The docked structure shows the non-basic face of PTN-NTD and the loop formed by residues 26 to 34 contact the α5-β5 and α6-β6 loop in α_M_I-domain (Figure 7). Besides hydrophobic contacts between L32 in PTN-NTD and I265 in α_M_I-domain as well as between PTN’s threonine methyls and P291 of α_M_I-domain, there were also several polar and electrostatic interactions in the interface, including between R261 in α_M_I-domain and D29 in PTN-NTD, R293 in α_M_I-domain and E36 in PTN-NTD (Figure 7).

**Figure 7.**
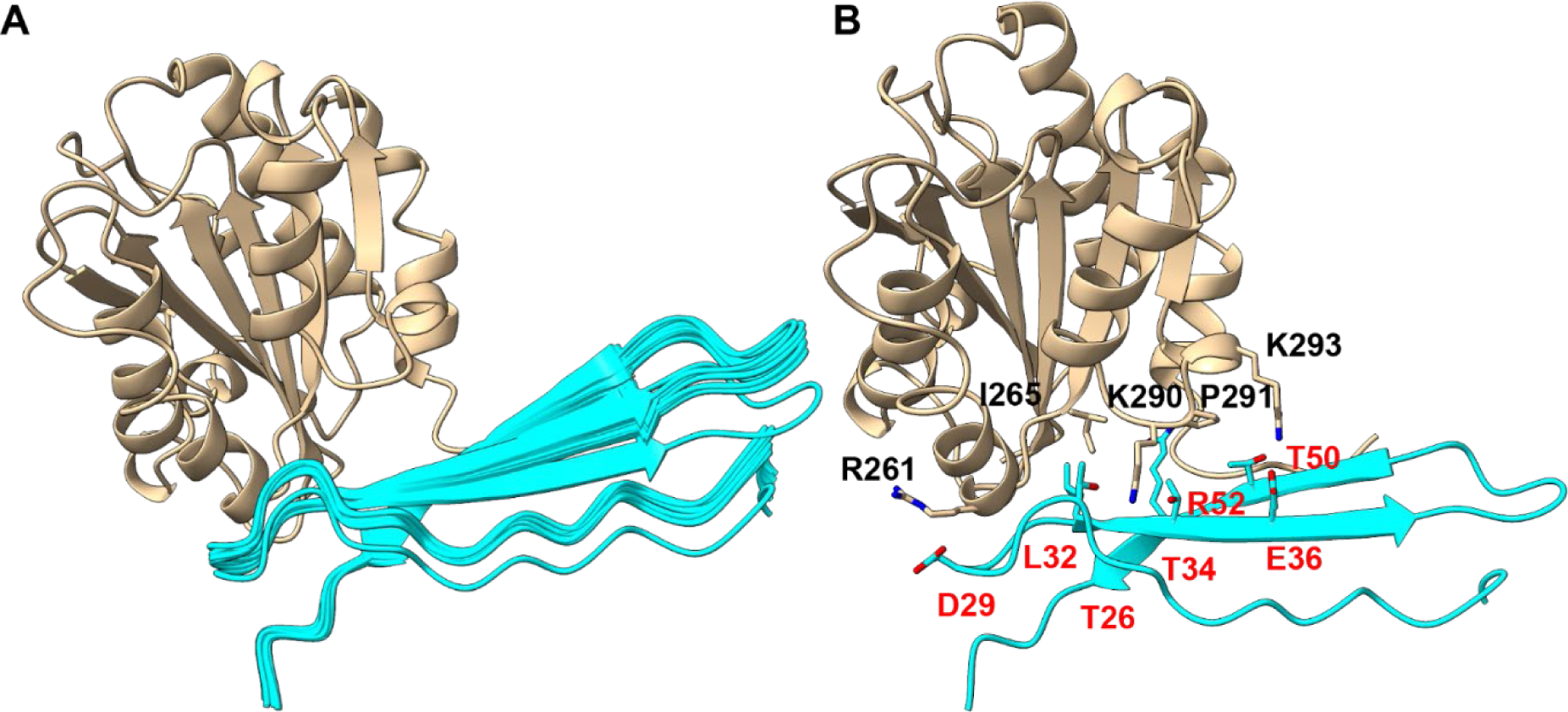
HADDOCK models of inactive α_M_I-domain bound to PTN-NTD. (A) Top 10 models with lowest HADDOCK scores after superimposing α_M_I-domain. The backbone RMSD of PTN-NTD is 1.9 Å. Only α_M_I-domain from model 1 is shown. (B) One of the structures from the best 10 structures with the lowest HADDOCK score showing contacts in the binding interface between the proteins.

We also docked PTN-CTD onto active α_M_I-domain using HADDOCK. A distance constraint between the MIDAS metal ion and the side chain of E98 was added based on crystal structures of active α_M_I-domain with other ligands (Bajic et al., 2013; Jensen et al., 2016; Trstenjak et al., 2020). In addition, we also added distance constraints extracted from the NOESY data (Table S5). In particular, distance constraints between residues A93 in PTN-CTD and residues S144 and R208 in active α_M_I-domain were used as well as constraints between E98 in PTN-CTD and residues G143, S144, I145, and R208 in active α_M_I-domain. HADDOCK clustered the 200 resulting structures into three clusters. Approximately 150 structures belonged to cluster 1. As expected, the main interaction is mediated by the 90s loop of PTN-CTD and MIDAS of α_M_I-domain. However, significant heterogeneity exists in the orientation PTN-CTD adopts relative to α_M_I-domain (Figure 8). As a result, the backbone RMSD for the structured portion of PTN-CTD (residues 66 to 109) after superimposing α_M_I-domain is 3.9 Å. Even though no distance constraints were specified between H95 of PTN-CTD and any residue in active α_M_I-domain, H95 was hydrogen bonded to the acidic amino acids in the acidic pocket formed by D242, E244, and D273 in active α_M_I-domain in some of the structures.

**Figure 8.**
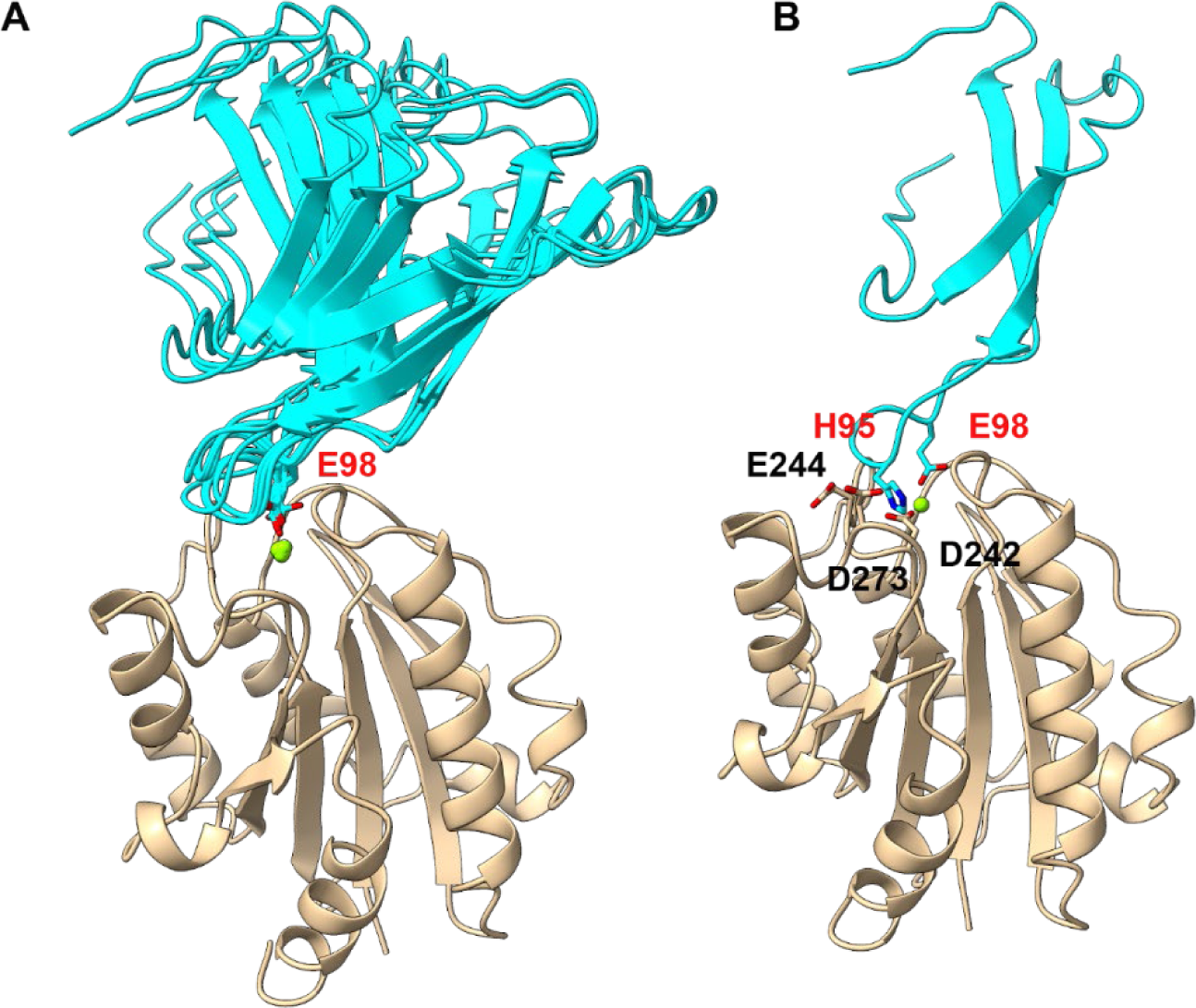
HADDOCK models of active α_M_I-domain bound to PTN-CTD. (A) Top 10 models with the lowest HADDOCK scores in cluster 1. The α_M_I-domain in the structures were superimposed. The backbone RMSD of PTN-CTD is 3.9 Å while the backbone RMSD of the binding loop (residues 92 to 101) is 2.5 Å. Only αMI-domain from model 1 is shown. (B) One of the structures with H95 close to the acidic pocket formed by D273, D242 and E244.

To investigate the stability and validity of the models, we carried out a 500-ns MD simulation of the complexes in explicit solvents using the software AMBER (Case et al., 2005). Figure 9A shows the RMSF of PTN-NTD backbone relative to the starting structure after superimposing the inactive α_M_I-domain backbone. Although a small shift in the position of the PTN-NTD at the beginning of the simulation was seen, PTN-NTD remained in stable contact with α_M_I-domain during the simulation. The RMSF of the PTN-NTD backbone for the last 100 ns of the simulation was only ∼ 2 Å (Figure 9A). In addition, the simulation revealed that the C-terminus of inactive α_M_I-domain can have significant electrostatic interactions with basic amino acids in PTN-NTD. In particular, residue E320 in inactive α_M_I-domain had strong interactions with both K49 and R52 in PTN-NTD, D294 in inactive α_M_I-domain hydrogen bonded to R52 in PTN-NTD, and the C-terminal carboxyl group of α_M_I-domain has interactions with K54 in PTN-NTD (Figure 9D, frame 1). The residues in the C-terminus of α_M_I-domain experienced significant PTN-induced chemical shift changes (Feng et al., 2021). However, NMR data showed no intermolecular NOEs between these residues and PTN-NTD. Results from the MD simulation indicate that the chemical shift perturbations may be due to dynamic electrostatic interactions between the C-terminus and the basic patch on PTN-NTD.

**Figure 9.**
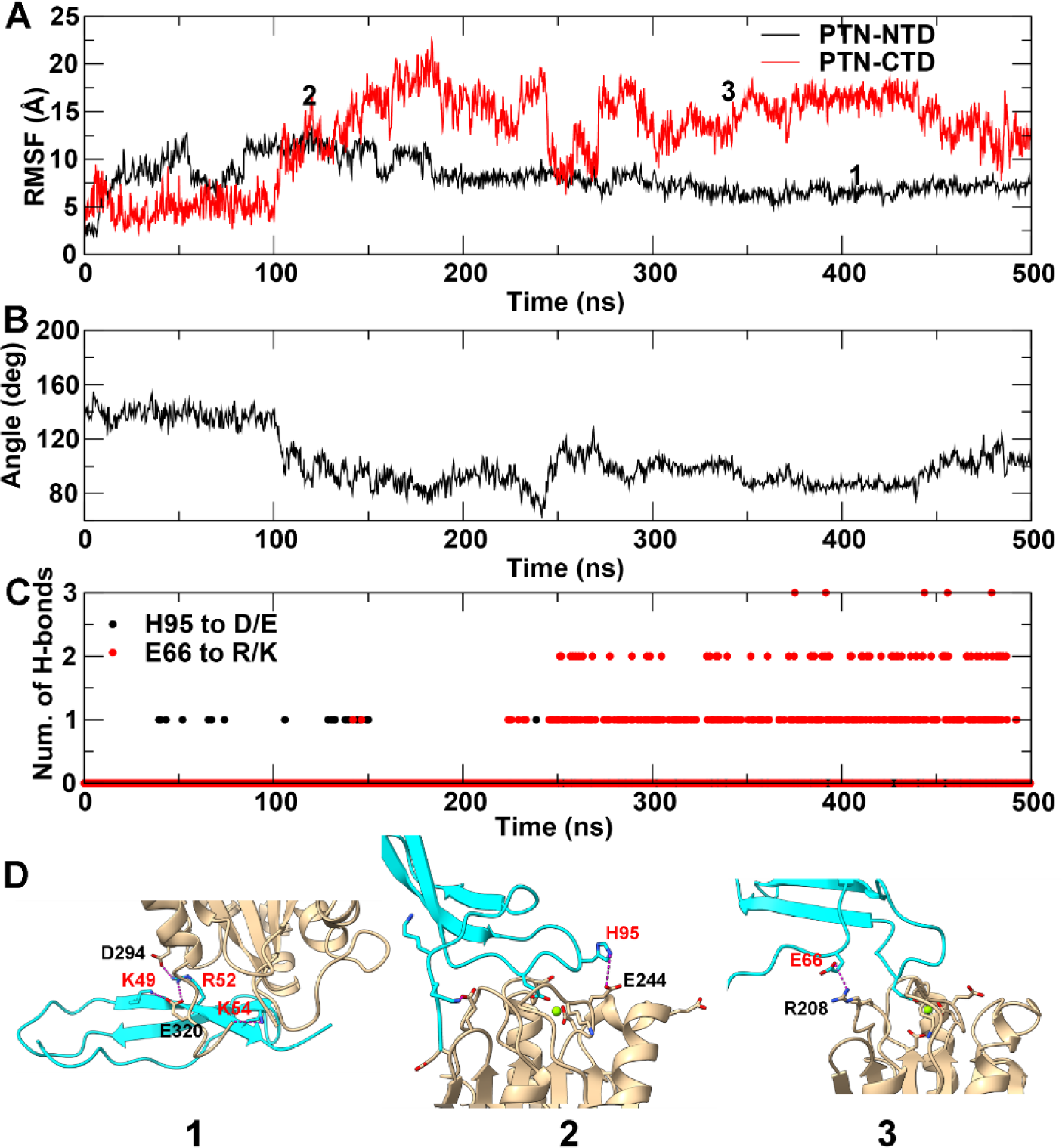
MD simulations of the models of PTN-NTD bound to inactive α_M_I-domain and PTN-CTD bound to active α_M_I-domain. A) RMSF of PTN-NTD and PTN-CTD structured regions relative to the starting structures after superimposing the α_M_I-domain. B) Changes in the orientation of PTN-CTD relative to α_M_I-domain during the simulation. The orientation is estimated by the angle between the vectors formed by the β1 strand in active α_M_I-domain (residues 133 to 140) and the middle β strand in PTN-CTD (residues 84 to 91). C) Intermolecular hydrogen bonds between H95 in PTN-CTD and Asp/Glu in α_M_I-domain (black), and E66 in PTN-CTD and Arg/Lys in α_M_I-domain (red) during the simulation of active α_M_I-domain bound to PTN-CTD. D) Frames from the simulations. 1. PTN-NTD’s basic face interacting with the C-terminus α_M_I-domain. 2. PTN-CTD’s H95 interacting with acidic amino acids near MIDAS. 3. PTN-CTD’s E66 interacting with R208 near the MIDAS. The positions of the frames in the simulations are marked with the frame numbers in panel A.

Similar MD simulations of PTN-CTD-bound to active α_M_I-domain were also carried out. We used a model with H95 in the acidic pocket as our starting structure. Similar to the HADDOCK results, PTN-CTD was considerably more dynamic than PTN-NTD in the simulation (Figure 9A). In particular, although the interaction of E98 with the MIDAS metal remained stable, the main body of PTN-CTD rotated by close to ∼35 ° before gaining stability (Figure 9B), causing H95 to to loss contact with the acidic pocket on the surface. However, the rotation allowed E66 of PTN-CTD to have electrostatic interactions with R208 in α_M_I-domain, compensating for the loss of H95’s interaction with α_M_I-domain (Figure 9C & 9D).

## DISCUSSION

In this study, we investigated the interaction of α_M_I-domain with PTN. Our data indicate that the α_M_I-domain can have both metal-dependent and metal-independent interactions with PTN. The metal-independent interaction is dominated by the binding of PTN-NTD to the bottom of α_M_I-domain and has fast time scale dynamics (Feng et al., 2021). This interaction also appears to be independent of the activation state of α_M_I-domain. The metal-mediated binding is between PTN-CTD and active α_M_I-domain. Its binding kinetics falls into the slow NMR time scale.

We previously reported that PTN-NTD is responsible for binding the inactive α_M_I-domain in a metal-independent fashion (Feng et al., 2021). In this study, we determined the high-resolution structure of the complex. The data revealed that, besides the α5-β5 loop, the α6-β6 loop of inactive α_M_I-domain also has extensive interactions with PTN-NTD. In particular, L32 of PTN-NTD contacts residues in the α5-β5 loop of inactive α_M_I-domain, with I265 being the most prominent residue in these interactions. Three threonines on one face of NTD also have extensive interactions with residues K290 and P291 in the α6-β6 loop. Finally, the side chain of R52 in PTN-NTD also contacted I265 from inactive α_M_I-domain. These contacts enabled us to model the structure of the complex using HADDOCK. Furthermore, MD simulations of the complex provided insight into why PTN perturbed the C-terminus of inactive α_M_I-domain. In particular, the simulation showed that the basic face of PTN-NTD has extensive electrostatic interactions with the C-terminus of α_M_I-domain. This explains the PTN-induced chemical shift perturbations previously observed in the C-terminus of inactive α_M_I-domain (Feng et al., 2021).

Besides the metal-independent interaction between inactive α_M_I-domain and PTN-NTD, our results indicate PTN-CTD can bind active α_M_I-domain using the canonical metal-chelation mechanism. Although many PTN-CTD acidic amino acids can chelate the divalent cation in the MIDAS, the most stable metal chelator in PTN is residue E98. PTN’s involvement in the metal-mediated binding mechanism is somewhat surprising because PTN’s highly positive net charge makes it an ideal basic ligand, many of whom are known to bind using a metal-free mechanism (Yakubenko et al., 2001). However, the few acidic amino acids in basic ligands may be sufficient to chelate the metal. In addition, even though PTN is highly basic, it does not have a significant amount of hydrophobic amino acids surrounding these basic amino acids, another feature required in the basic protein binding motif (Podolnikova et al., 2015b). Therefore, it may be unable to take advantage of α_M_I-domain’s binding site for basic/hydrophobic ligands.

One interesting finding of the study is that no single acidic amino acid in PTN is essential to binding. This implies that active α_M_I-domain does not chelate just one acidic amino acid in PTN but can interact with multiple acidic amino acids. Signs of heterogeneous binding modes were also reported for the interaction of denatured fibrinogen with active α_M_I-domain as well as α_X_I-domain (Vorup-Jensen et al., 2005). LL-37’s interaction with active α_M_I-domain also appears to be heterogeneous in SPR analysis (Zhang et al., 2016). The heterogeneity explains why the PCS and non-PCS signals coexist and the intensities of PCS signals are only about 11 % of the normal peak. In particular, this may reflect that PCS signals can only be produced when E98 is the metal chelator and active α_M_I-domain doesn’t always bind E98. E98’s ability to produce PCS is most likely the result of other interactions stabilizing PTN’s interaction with α_M_I-domain when E98 is in the MIDAS. One contact that may contribute to this is the interaction between PTN’s H95 and the acidic pocket near MIDAS formed by D242, E244, and D273. The importance of H95 was demonstrated by the fact that its mutation to serine produced significant changes in PTN-CTD induced chemical shift perturbations observed in the ^15^N-HSQC spectrum of α_M_I-domain. Mutating H95 to anything other than another basic amino acid also resulted in the loss of all PCS. Chelation of the MIDAS metal by other PTN acidic amino acids besides E98 makes this interaction impossible and can result in dynamics that average the PCS to zero. However, the small magnitude of PCS observed in PTN indicates that even the interaction mediated by E98 chelation may be dynamic. This agrees with HADDOCK modeling and MD simulation, both of which showed dynamic movements in PTN-CTD’s interaction with active α_M_I-domain when E98 is the chelator. It is worth noting that conformational dynamics were also observed in the crystal structure of the drug simvastatin bound to active α_M_I-domain (Jensen et al., 2016).

H95 and E98 can also be viewed as forming a zwitterionic ligand. In this regard, the interaction of active α_M_I-domain with PTN is akin to integrins that bind the zwitterionic RGD motif across two different domains in the α and β subunits. However, in the case of α_M_I-domain, the binding sites for the positive and negative ions are found on the same domain. In addition, α_M_I-domain is not the only α I-domain with a preference for zwitterionic ligands. A homologous acidic pocket in the human α_L_I-domain was shown to be crucial to the binding of the zwitterionic motif ^34^ETPLPK^39^ in ICAM-1 (Shimaoka et al., 2003). The same phenomenon likely exists in ICAM-1 and α_M_I-domain binding. In particular, it has been proposed that D229 in D3 of ICAM-1 is most likely the metal chelator (Diamond et al., 1991; Mao et al., 2011). Interestingly, R231 is situated nearby and the side chains of D229 and R231 point in the same direction, enabling R231 to bind the acidic pocket of D242, E244, and D273 on α_M_I-domain. Such zwitterionic interaction was also observed in leukocidin GH’s interaction with α_M_I-domain. In particular, R294 and K319 in leukocidin H bind the acidic pocket while E323 of leukocidin H chelates the metal in α_M_I-domain (Trstenjak et al., 2020). The same phenomenon was also proposed for GP1bα’s interaction with α_M_I-domain with H220 in GP1bα playing the role of the basic amino acid while E224 chelates the metal (Morgan et al., 2019). In addition, the removal of K39 in ICAM-1, K319 in leukocidin H, and H220 in GP1bα significantly reduced the binding of the respective ligand to α_M_I-domain (Shimaoka et al., 2003; Morgan et al., 2019; Trstenjak et al., 2020). However, in the case of PTN, removing H95 only eliminated PCS experienced by PTN without changing the binding affinity. This suggests that H95’s interaction with the acidic pocket may not be as strong as in other ligands. Factors such as the accessibility of E98, and the favorable interaction between residue E66 in PTN and residue R208 in α_M_I-domain, may also contribute to α_M_I-domain’s preference for E98. It should be noted that other ligands also utilize residue R208 in their interaction with α_M_I-domain. In particular, acidic amino acids in both C3d and leukocidin H have contacts with residue R208 in α_M_I-domain. C3d also has electrostatic interactions with residues E178 and E179 located next to R208 (Bajic et al., 2013; Trstenjak et al., 2020). The emerging trend from these studies is that the charged residues around α_M_I-domain’s MIDAS play important roles in mediating interaction with ligands.

It should also be noted that the acidic pocket next to the MIDAS may serve as a ligand-binding site independent of the MIDAS. In particular, the acidic pocket can be an ideal binding site for basic proteins and peptides, which are known to have strong affinities for active α_M_I-domain (Podolnikova et al., 2015b; Lishko et al., 2018). Although the binding site for most of the basic ligands has not been confirmed, a previous study on the interaction between α_M_I-domain and the archetypal basic α_M_I-domain ligand, the peptide P2-C from fibrinogen, showed that mutations of residues around MIDAS significantly attenuated the binding of P2-C to α_M_I-domain (Yakubenko et al., 2001). This strongly supports the proposal that basic ligands can bind to sites around MIDAS.

The finding that both PTN domains are involved in α_M_I-domain binding suggests a mechanism by which PTN may cross-link cells to the extracellular matrix. In particular, because both domains of PTN can bind GAG as well as active α_M_I-domain, it is plausible that one domain may bind GAG while the other binds α_M_I-domain. It is also tempting to speculate whether NTD and CTD from the same molecule of PTN can simultaneously bind the MIDAS and N/C-termini sites. However, the short linker between NTD and CTD makes such a scenario sterically challenging. The finding that wild-type PTN’s affinity for active α_M_I-domain is no higher than that of PTN-CTD also supports the lack of simultaneous binding of α_M_I-domain by both PTN domains from the same PTN molecule. In addition, PTN’s binding site for active α_M_I-domain does not overlap with PTN’s GAG-binding site. This explains why a previous study has shown that PTN immobilized on proteoglycans can still support macrophage adhesion (Shen et al., 2017)

The electrostatic surface potentials of α_L_I-domain and α_X_I-domain are significantly different from that of α_M_I-domain (Vorup-Jensen and Jensen, 2018). In particular, the α_L_I-domain has a hydrophobic patch near its MIDAS in addition to the acidic pocket. This patch is absent in both α_M_I-domain and α_X_I-domain and may be the reason behind α_L_I-domain’s monospecificity. In particular, the hydrophobic patch helps to exclude solvent from the MIDAS of α_L_I-domain, thereby strengthening the electrostatic interaction between the ligand and the divalent cation. This may be the reason behind α_L_I-domain’s high affinity for domain 1 of ICAM-1 (Shimaoka et al., 2003; San Sebastian et al., 2006). Similar hydrophobic interaction with other ligands may be a prerequisite for achieving strong affinity for α_L_I-domain. The lack of this hydrophobic patch in α_M_I-domain and α_X_I-domain suggests that these domains may be less selective and would bind any charged ligands, albeit at lower affinity. This feature may partly explain ligand binding promiscuity exhibited by Mac-1. The ligand specificities of α_M_I-domain and α_X_I-domain are also not identical. The basis for this difference lie in the absence of the acidic pocket in α_X_I-domain. The lack of an acidic pocket near the MIDAS significantly enhances α_X_I-domain’s affinity for anionic polymers such as heparin and unfolded proteins compared to α_M_I-domain or α_L_I-domain (Vorup-Jensen et al., 2005). These differences may be the key to developing specific inhibitors for each α I-domain.

In summary, we have determined the interaction between PTN and α_M_I-domain. We conclude that PTN can bind α_M_I-domain using two different mechanisms depending on the activation state of α_M_I-domain. When α_M_I-domain is in the inactive state, PTN binds to the bottom side of α_M_I-domain using PTN-NTD and a metal-independent mechanism. When α_M_I-domain is in the active state, the interaction is dominated by the canonical metal-chelation mechanism in which PTN’s residue E98 acts as the major chelator of the divalent cation in the MIDAS. In addition, the chelation of the metal by E98 is stabilized by favorable electrostatic interactions between PTN and active α_M_I-domain residues near the MIDAS. We think these interactions are crucial to determining the ligand specificity of α_M_I-domain.

## KEY RESOURCES TABLE

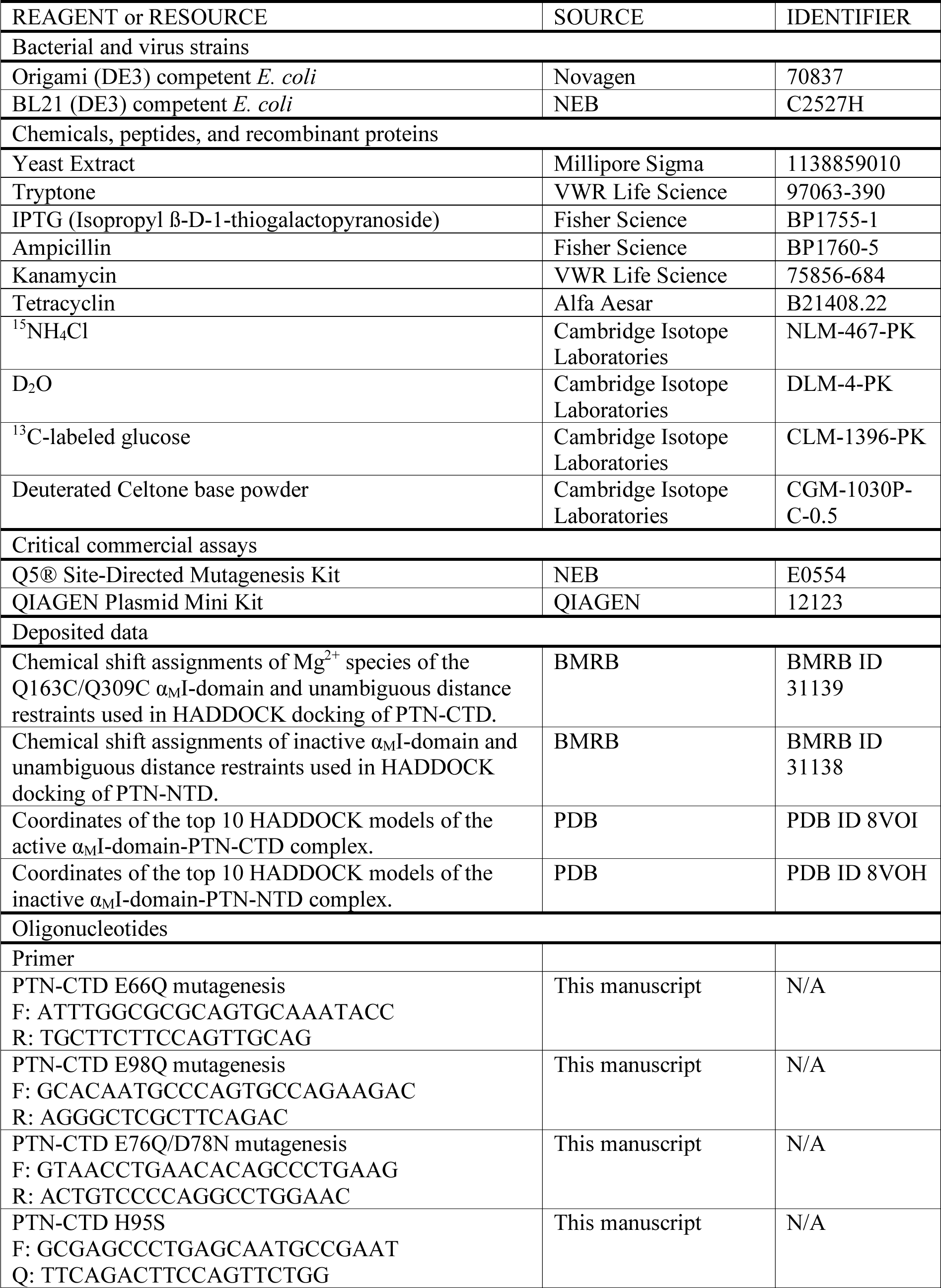
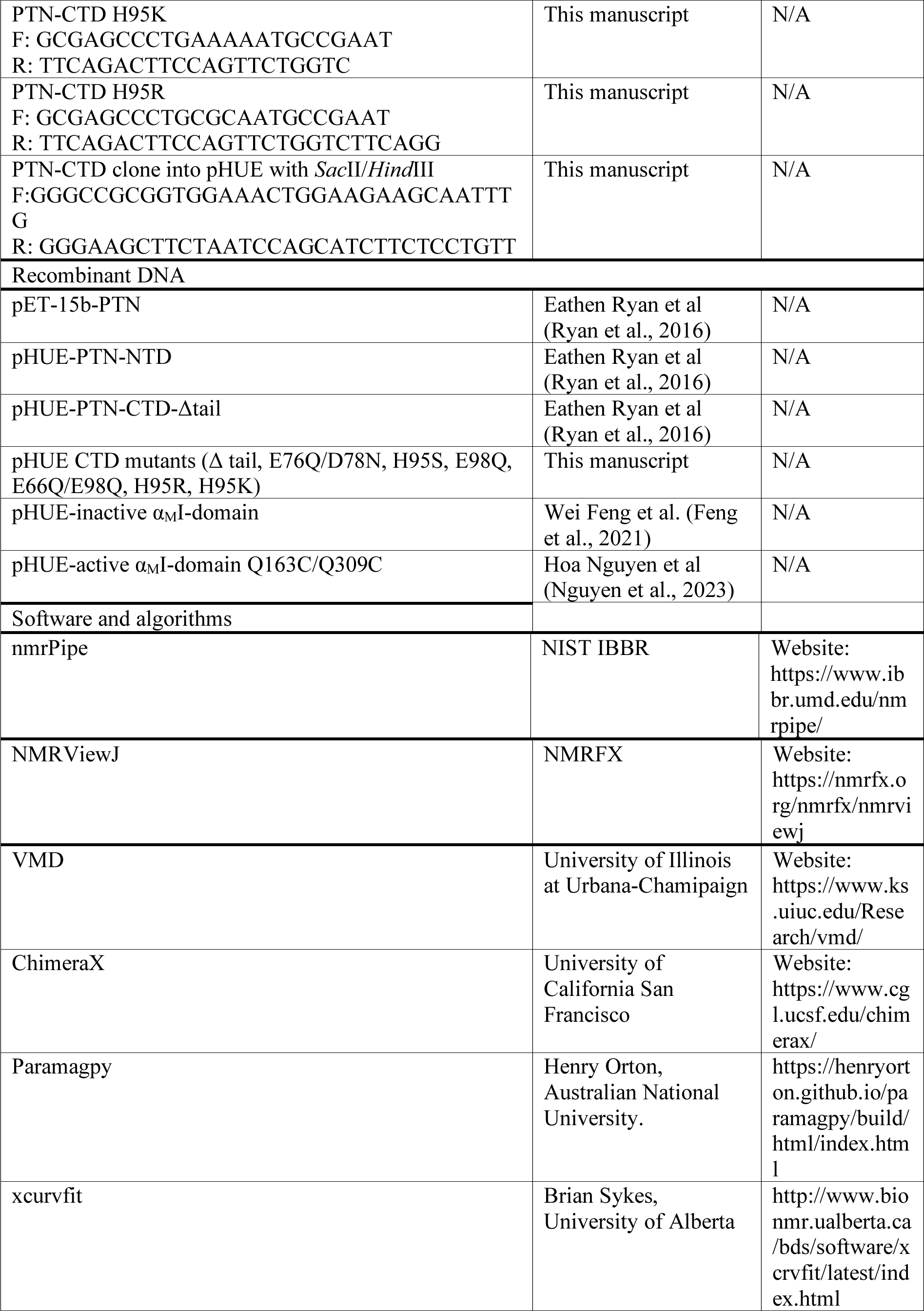
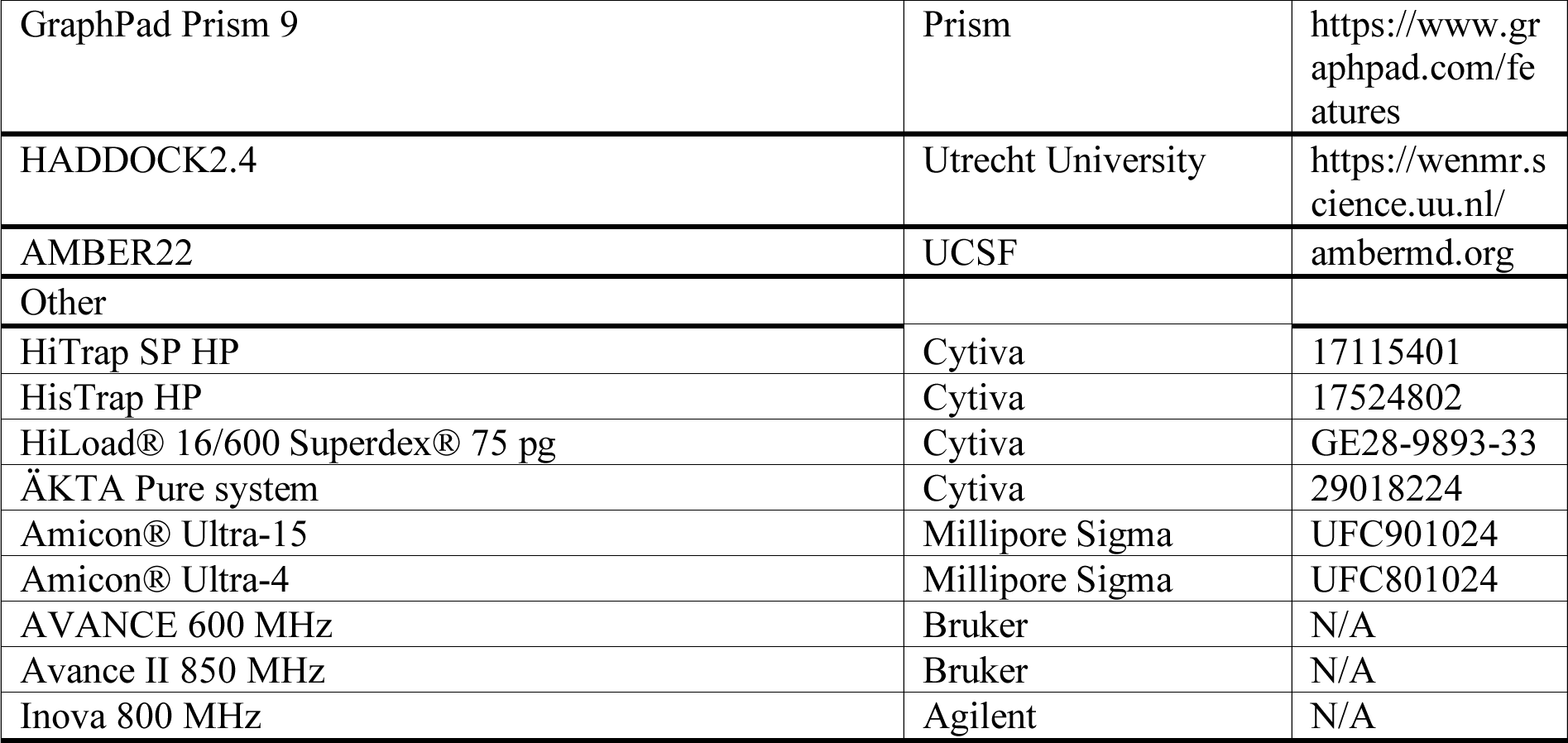

## METHOD DETAILS

### Expression and Purification of α_M_I-domains

The expression and purification of α_M_I-domain were accomplished using the previously reported procedures (Feng et al., 2021; Nguyen et al., 2023). Briefly, the open reading frame (ORF) of the wild-type human α_M_I-domain (E131-T324) was cloned into the pHUE vector (Catanzariti et al., 2004) using *Sac*II and *Hind*III as restriction sites. The activating Q163C/Q309C cysteine mutations were introduced into α_M_I-domain using the Q5 Site-Directed Mutagenesis Kit (NEB). The plasmids were transformed into either BL21(DE3) (NEB) or OrigamiB(DE3) (Novagen) and the cells were grown in M9 media at 37 ^°^C until the culture reached an OD_600_ of ∼ 0.8-1. The protein expression was induced with 0.5 mM IPTG, and the cell pellet was harvested after overnight incubation at 22°C. The pellet was resuspended and incubated on ice in the lysis buffer (20 mM sodium phosphate, 0.5 M NaCl, 10 mM imidazole, 5% glycerol) containing 1 mg/ml lysozyme for 20 min on ice. After sonication and centrifugation, the supernatant was loaded onto a 5-mL HisTrap column (Cytiva), and the protein was eluted from the column by running a 0.01 to 0.5 M imidazole gradient. To separate His-ubiquitin from α_M_I-domain, the protein was digested with enzyme USP2 (1:50 molar ratio) overnight at room temperature in 20 mM Tris, 0.1 M NaCl (Catanzariti et al., 2004). The digestion mixture was subjected to Ni^2+^ column purification again. α_M_I-domain in the flow-through was further purified with a 120-mL Superdex 75 column using a buffer containing 20 mM HEPES, 0.3 M NaCl, pH 7.0. Finally, the protein was exchanged into 20 mM HEPES, 0.1 M NaCl, pH 7.0 for NMR analyses. To prepare isotope labeled protein, ^15^N, ^13^C or ^2^H labeling was accomplished by adding ^15^NH_4_Cl, ^13^C glucose, D_2_O, and deuterated Celtone base powder (Cambridge Isotope). The ^2^H, ^15^N-labeled α_M_I-domain was prepared by seeding freshly transformed colony in 20 mL of LB until the OD_600_ reached 1.0. Cells from 5 mL of the LB culture were gently pelleted, used to inoculate 50 mL of ^2^H, ^15^N minimal media, and grew overnight at 37 °C. The 50-mL culture was then used to seed 0.45 L of media the next day and grown as described before. The protein was expressed and harvested with the same procedure mentioned above.

### Expression and Purification of PTN

PTN and its domains were produced following previously reported protocols (Feng et al., 2021). Briefly, the mature PTN open reading frame was cloned into a pET-15b vector using *Nco*I and *Xho*I restriction sites and transformed into OrigamiB(DE3) cells. The transformed cells were grown under conditions similar to the Q163C/Q309C mutant of the α_M_I-domain. To extract the protein, cell pellets were resuspended in 20 mM Tris, 0.2 M NaCl, pH 8.0, 1 mg/mL of lysozyme and incubated for 20 minutes at room temperature. After sonication and centrifugation to remove insoluble material, the protein is extracted from the supernatant using a 5-mL HiTrap SP HP column (Cytiva) and eluted with a 0.1-1.5 M NaCl gradient. To produce PTN domains, a pHUE vector(Catanzariti et al., 2004) containing PTN ORF coding either residues G1 to C57 (PTN-NTD), residues N58 to K114 (PTN-CTD Δtail), or residues N58 to D136 (PTN-CTD) at the 3’ end of the ubiquitin ORF was transformed into OrigamiB(DE3) cells. Cells were grown in M9 media at 37 °C to an OD_600_ of 0.8. 0.25 mM IPTG was added to the culture and incubated overnight at 23°C. Purifications of PTN domains and various mutants were similar to that of α_M_I-domain except no size exclusion chromatography was used.

### NMR Data Acquisition

NMR data were collected on either Bruker AVANCE 600 MHz equipped with a Prodigy probe, a Bruker Avance II 850 MHz equipped with a cryoprobe, or an Agilent Inova 800 MHz equipped with a cold probe. All data were collected at 25 °C. Backbone assignments of PTN and Q163C/Q309C α_M_I-domain were carried out previously (Ryan et al., 2016; Nguyen et al., 2023). For PTN domain backbone and side chain assignment, CACBCONH, HNCACB, and HCCCONH were collected on ^13^C, ^15^N-labeled PTN-NTD and PTN-CTD. The parameter was set with 1024 complex points in ^1^H dimension, 30 complex points in ^15^N dimension, 50 complex points in ^13^C dimension for 3D experiment. 2D ^15^N-HSQC experiments were acquired with 1024 complex points in ^1^H dimension and 50 complex points in the ^15^N dimension with the spectrum widths are 15 ppm for ^1^H dimension and 35 ppm for ^15^N dimension with carrier at 4.7 and 119ppm, respectively.

Intermolecular NOEs between PTN-NTD and inactive α_M_I-domain were obtained using F1-^13^C-edited/F3-^13^C,^15^N-filtered HSQCNOESY experiments. The sample used to collect these data contained 0.2 mM ^13^C-labeled inactive α_M_I-domain and 1 mM unlabeled PTN-NTD. To confirm the assignments of PTN-NTD atoms at the interface, we acquired F1-^13^C,^15^N-filtered /F3-^13^C-edited NOESYHSQC data on samples containing 0.2 mM unlabeled inactive α_M_I-domain and 0.5 mM ^13^C, ^15^N-labeled PTN-NTD. To obtain intermolecular NOE between PTN-CTD and active α_M_I-domain, we collected 2D projections of the ^13^C HSQC - NOESY-^15^N HMQC experiment on a sample containing 0.25 mM of ^2^H,^15^N-labeled Q163C/Q309C α_M_I-domain and 1 mM ^13^C PTN-CTD in 20 mM HEPES, 0.1 M NaCl, pH 7.0. The NOE mixing time is 0.2s.

Each titration of active α_M_I-domain with PTN mutants was performed with a series of six samples. All samples in the series contained ∼ 90 µM of ^15^N-labeled active α_M_I-domain. The PTN ligand concentrations in the samples are 0, 0.3 0.6, 0.9, 1.2 and 1.5 mM. ^15^N-HSQC of each sample was acquired with the parameters described above.

To measure the α_M_I-domain-induced chemical shift changes on PTN-NTD, ^15^N HSQC spectra were acquired on a sample containing 0.1 mM of ^15^N-labeled PTN-NTD and 0.8 mM of unlabeled inactive α_M_I-domain or Q163C/Q309C α_M_I-domain. The sample buffer is 20 mM HEPES, 0.1 M NaCl, pH 7.0. The chemical shift change of each signal was quantified using the equation δ=[Δδ_H2_ + (0.17 Δδ_N_)^2^]^1/2^, where Δδ_H_ is the chemical shift change of amide hydrogen and Δδ_N_ is the chemical shift change of amide nitrogen.

To confirm that PTN does not induce significant changes in the structure of inactive α_M_I-domain, an aligned sample of 0.1 mM ^15^N α_M_I-domain in a 6% neutral polyacrylamide gel (Cierpicki and Bushweller, 2004) was used to obtain HN residual dipolar coupling (RDC). The RDC of α_M_I-domain were measured in the presence and absence of 0.8 mM unlabeled PTN-NTD.

To measure the pseudocontact shift (PCS) of α_M_I-domain in the presence and absence of PTN-NTD and PTN-CTD, we collected ^15^N-HSQC, HNCACB and CBCACONH spectra of Co^2+^-saturated inactive α_M_I-domain in the presence of unlabeled PTN-NTD and of Co^2+^-saturated active α_M_I-domain in the presence of unlabeled PTN-CTD. The NMR samples used to collect these data contained 0.2 mM ^13^C, ^15^N α_M_I-domain, 1 mM unlabeled PTN-CTD or PTN-NTD, 2 mM of either CoCl_2_ or MgCl_2_, 20 mM HEPES, 0.1 M NaCl, pH 7.0. The amide hydrogen and nitrogen chemical shift assignments of α_M_I-domain residues in the presence of Co^2+^ were obtained by comparing H, N, CA, CB chemical shifts of each spin system with the corresponding chemical shifts from residues in the Mg^2+^ sample. A match is made if the four chemical shifts from a spin system in the Co^2+^ data match the four chemical shifts from an assigned residue in the Mg^2+^ sample while taking into consideration the PCS on each atom as a result of the Co^2+^. PCS tensors were calculated using the software Paramagpy (Orton et al., 2020) using only the PCS measured from amide hydrogens and PDB structures 1JLM (inactive α_M_I-domain) or 1IDO (active α_M_I-domain).

### Modeling of PTN-α_M_I-domain complexes

The model between α_M_I-domain and PTN domains was generated using HADDOCK (Dominguez et al., 2003). Unambiguous contacts between α_M_I-domain and PTN-NTD’s residues obtained from F1-^13^C-edited/F3-^13^C,^15^N-filtered HSQCNOESY spectra were used as constraints for the model (Table S4). For the complex between active α_M_I-domain and PTN-CTD, contacts observed in the 4D ^13^C-HSQC-NOESY-^15^N-HMQC spectrum were used as constraints (Table S5). The crystal structure of inactive α_M_I-domain (PDB accession number 1JLM) and the solution structure of PTN-NTD (PDB accession number 2N6F) were used as the starting structures to model the Mg^2+^-independent interactions between α_M_I-domain and PTN-NTD. The crystal structure of the active α_M_I-domain (PDB accession number 1IDO) was used as the starting α_M_I-domain structure to model the Mg^2+^-dependent interaction. During docking, we allowed the protein segments containing residues involved in binding to be fully flexible. For the modeling of Mg^2+^-independent interaction, these residues include D260 to R266 and I287 to V296 from α_M_I-domain, as well as P25 to S27, L32 to R34, and Q46 to R52 from PTN-NTD. For the modeling of Mg^2+^-dependent interaction, flexible residues include G141 to I145 and L206 to R208 in α_M_I-domain, and A93 to E98 in PTN-CTD. The docking followed the default protocol with an explicit solvent refinement for the last cycle. The best models were chosen based on HADDOCK energy scores and agreement with the constraints. Molecular dynamics (MD) simulations of the models were carried out using the top-scoring model from each docking study as the starting structure. For each simulation, the starting structure was first energy minimized. This was followed by a gradual heating to 300 K and then equilibration at 300 K with a 50 ps simulation. The production run was conducted with constant pressure and simulated with 1 fs step for 500 ns.

### Resource Availability

#### Lead contact

Further information and requests for resources and reagents should be directed to and will be fulfilled by Xu Wang.

#### Materials availability

All unique reagents generated in this study are available from the lead contact without restriction.

#### Data availability

HADDOCK models of the inactive α_M_I-domain-PTN-NTD complex have been deposited as PDB entry 8VOH. HADDOCK models of the active α_M_I-domain-PTN-CTD complex have been deposited as 8VOI. Restraints and chemical shifts used in the modeling have been deposited as BMRB entry 31138 for the inactive α_M_I-domain-PTN-NTD complex and as BRMB entry 31139 for the active α_M_I-domain-PTN-CTD complex.

## ACKNOWLEDGEMENTS

We thank the staff of the Magnetic Resonance Research Center at Arizona State University for the maintenance of the NMR instruments. The study was funded by NIH grants R01GM118518 (to X.W.) and R01HL063199 (to T.U.).

## Author Contribution

X.W. and T. U. conceived the project. X. W. and H. N. designed and conducted the experiments. X. W., H.N., N. P., and T. U. wrote the paper.

## Declaration of interests

The authors have no competing interests.

## Notes

### Competing Interest Statement

The authors have declared no competing interest.

