## Supplementary file for "*α*_M_I-domain of Integrin Mac-1 Binds the Cytokine Pleiotrophin Using Multiple Mechanisms"

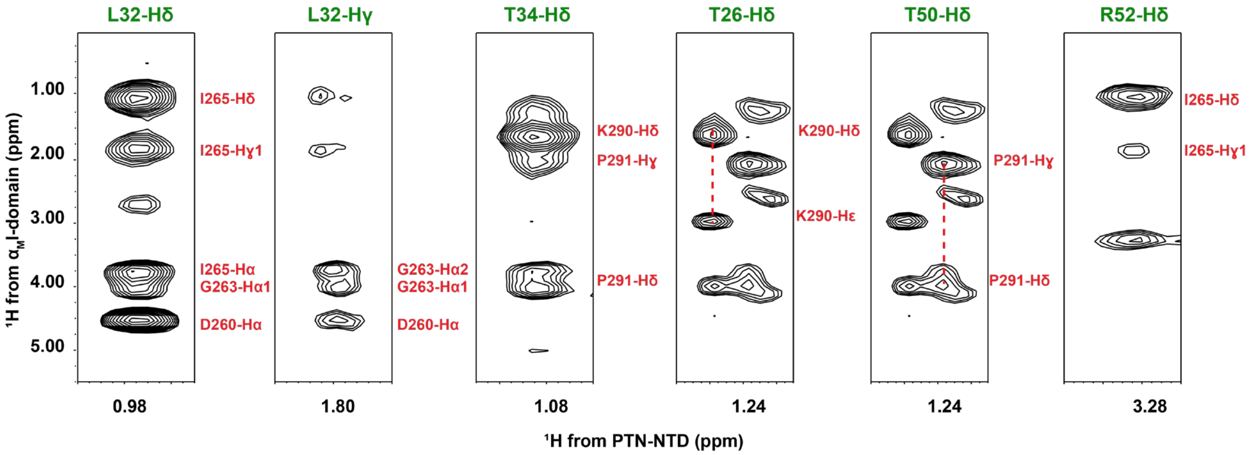


**Figure S1**. F1-$^{13}$C,$^{15}$N-filtered /F3-$^{13}$C-edited NOESYHSQC spectrum of 0.2 mM unlabeled $\alpha_M$I-domain in the presence of 0.5 mM $^{13}$C,$^{15}$N-labeled PTN-NTD. The experiment allows us to obtain the $^{13}$C chemical shifts of PTN-NTD residues involved in the interaction with inactive $\alpha_M$I-domain.

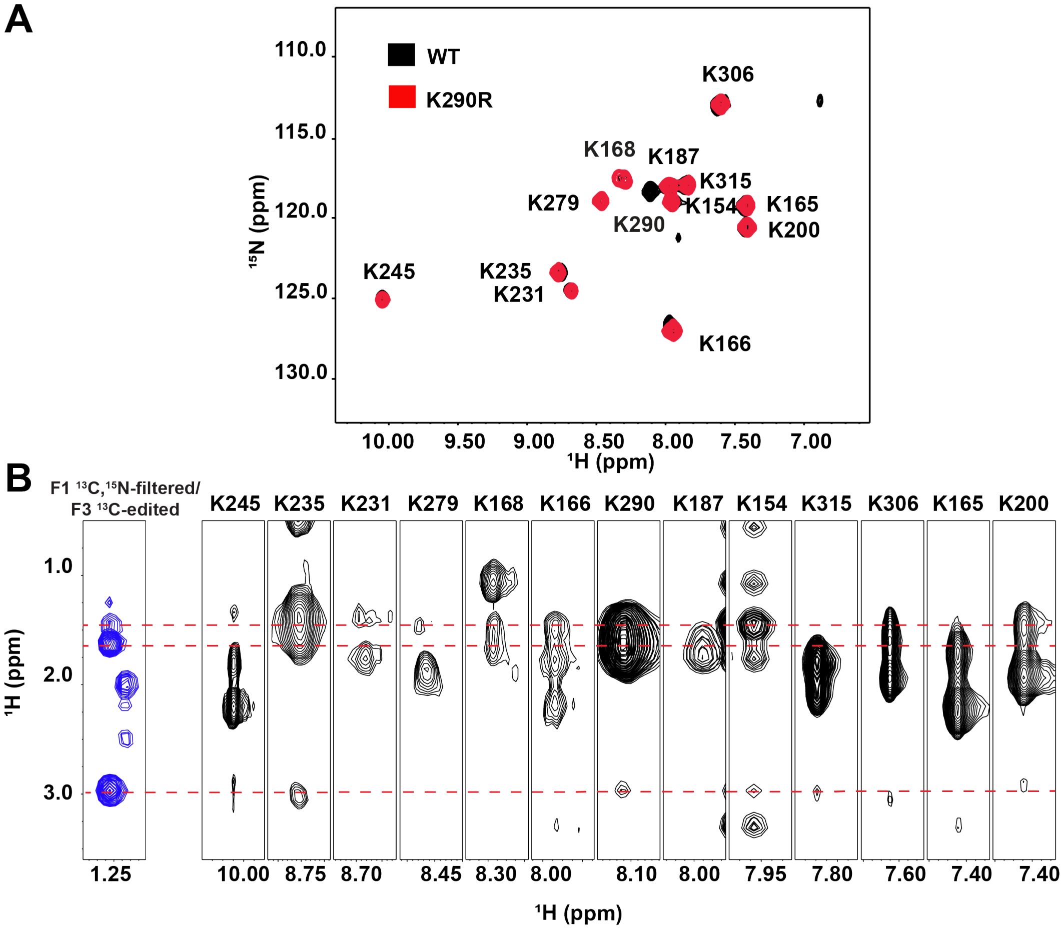


**Figure S2**. (A) The assignment of residue K290 in $\alpha_M$I-domain was confirmed through selective $^{15}$N-labeling of lysines of inactive $\alpha_M$I-domain (black) and the K290R mutant (red). B) Patterns of intermolecular NOE between $\alpha_M$I-domain and PTN-NTD compared with intramolecular NOE patterns of $\alpha_M$I-domain lysines. Only K290 proton side chain chemical shifts match the intermolecular NOE cross peaks seen in the F1-$^{13}$C-edited/F3-$^{13}$C,$^{15}$N- filtered HSQCNOESY experiments.

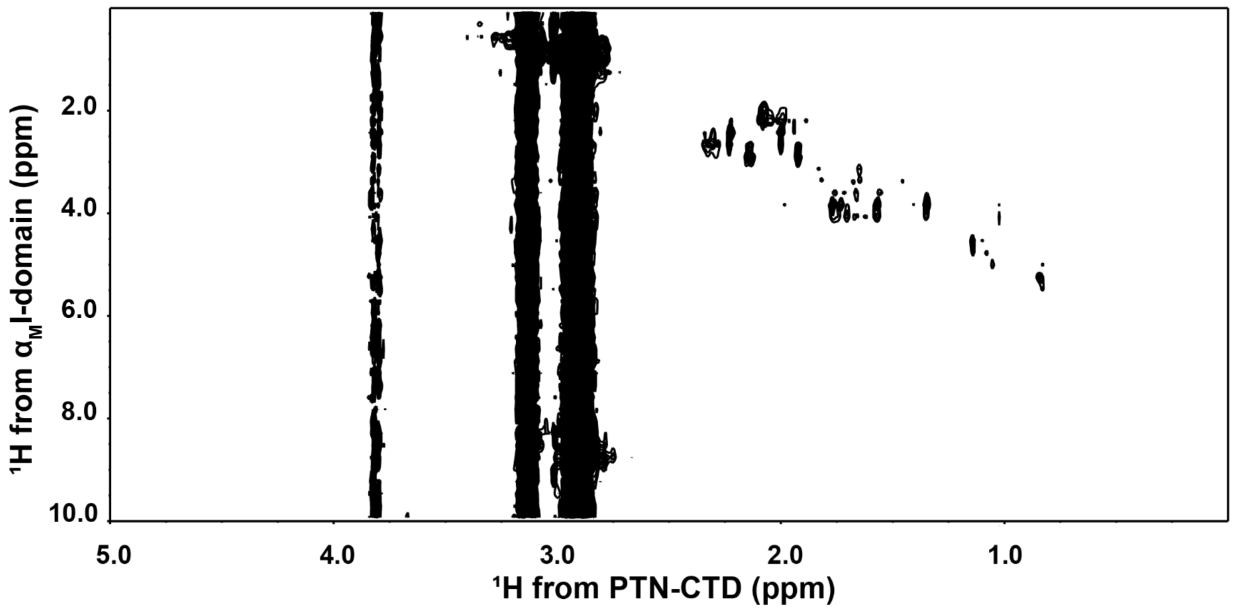


**Figure S3**. Projection of F1-$^{13}C$-edited/F3-$^{13}C$,$^{15}N$-filtered HSQCNOESY spectrum of 0.2 mM $^{13}C$,$^{15}N$-labeled inactive $\alpha_M$I-domain in the presence of 0.7 mM unlabeled PTN-CTD. The data produced no identifiable intermolecular cross peaks between PTN-CTD and inactive $\alpha_M$I-domain. The cross peaks are the result of break through diagonal peaks.

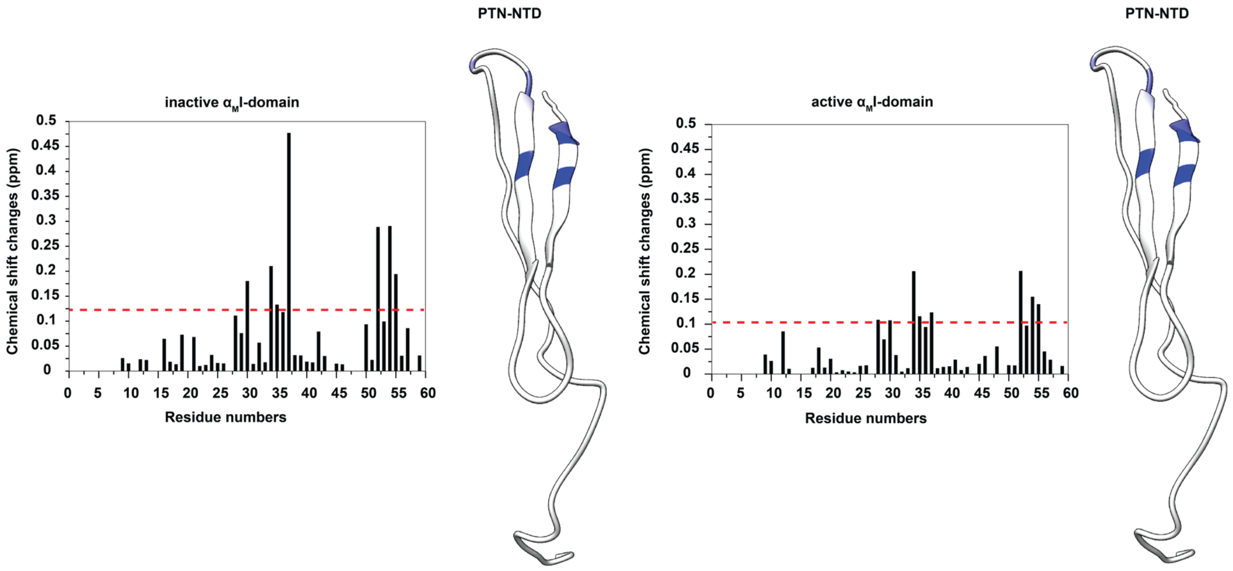


**Figure S4**. Residue specific chemical shift changes in 0.1 mM $^{15}$N PTN-NTD induced by 0.6 mM unlabeled inactive and active $\alpha_M$I-domain. The value of 1.5 standard deviations higher than the average chemical shift change of all residues is indicated by the red line. The ribbon representation of the PTN-NTD with residues showing $\alpha_M$I-domain-induced chemical shift change greater than 1.5 SD above average are shown in blue. Inactive and active $\alpha_M$I-domains produced similar chemical shift changes in PTN-NTD, indicating that PTN-NTD's interaction with the $\alpha_M$I-domains is independent on its activation state.

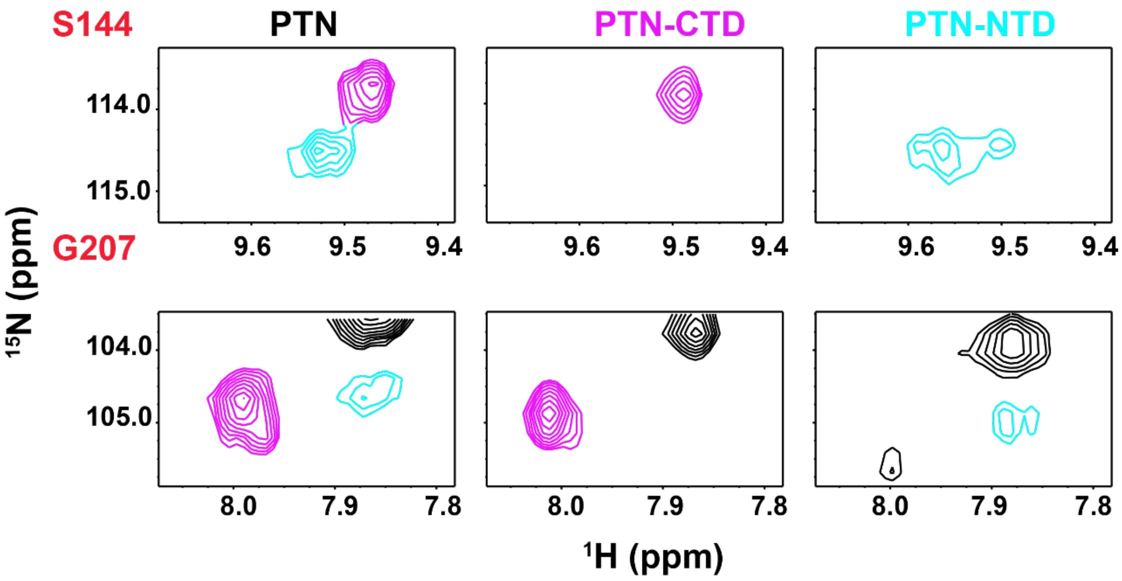


**Figure S5**. Heterogeneity in PTN-induced changes in the $^{15}$N-HSQC signals of $\alpha_M$I-domain's MIDAS residues can be seen in sections of $^{15}$N-HSQCs of active $\alpha_M$I-domain in the presence of wild type PTN (black) or PTN domains (CTD-magenta, NTD-cyan). Binding of PTN domains produces domain specific signals in some MIDAS residue signals, such as G143, S144 and G207.

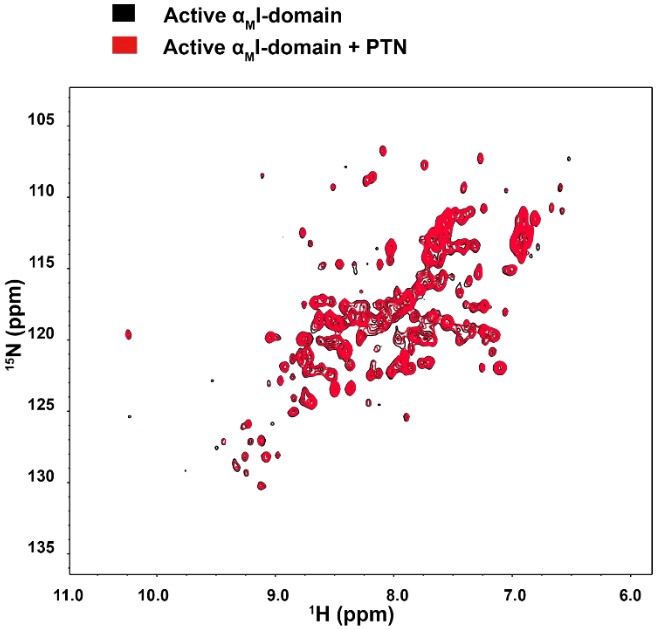


**Figure S6**. Binding of PTN requires $Mg^{2+}$. Spectra of apo active $\alpha_M$I-domain with and without PTN. Spectral changes indicative of MIDAS residue perturbations were not seen.

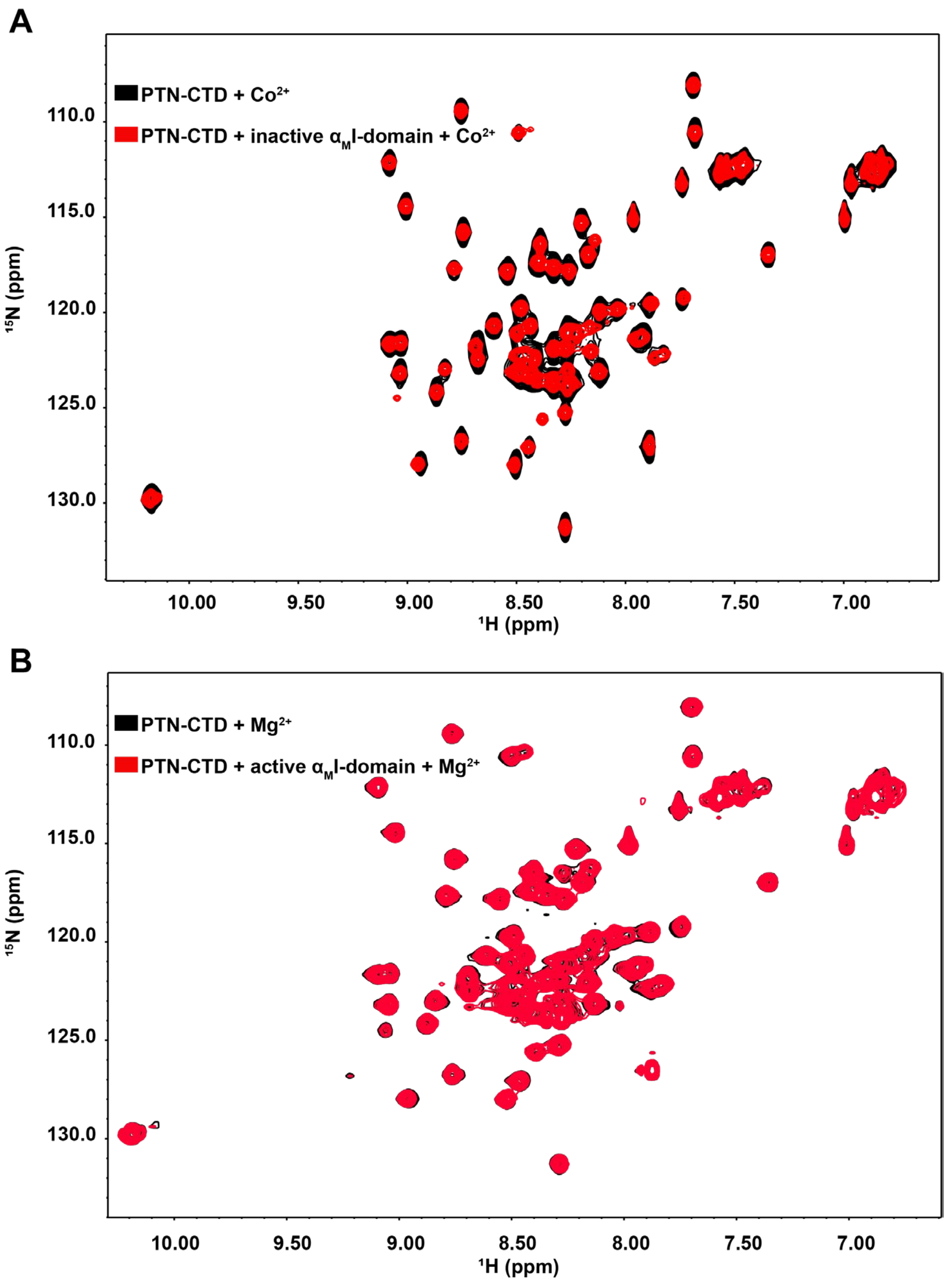


**Figure S7**. (A) $^{15}$N-HSQC spectrum of 0.1 mM $^{15}$N PTN-CTD with $Co^{2+}$ and in the absence (black) or presence (red) of 0.2 mM unlabeled inactive $\alpha_M$I-domain. (B) $^{15}$N-HSQC spectrum of 0.1 mM $^{15}$N PTN-CTD with $Mg^{2+}$ and in the absence (black) or presence of (red) of 0.2 mM unlabeled active $\alpha_M$I-domain. In both case, PCS signals are not induced by either $Co^{2+}$ or active $\alpha_M$I-domain alone.

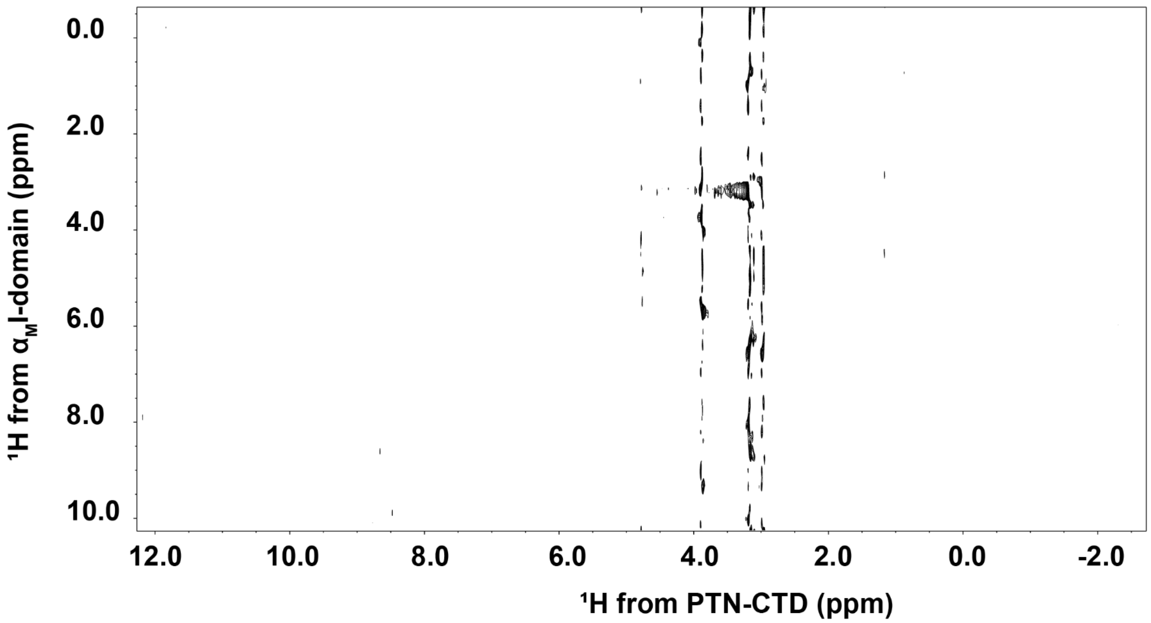


**Figure S8**. $^1H$-$^1H$ projection of 3D F1-$^{13}C$-edited/F3-$^{13}C$,$^{15}N$-filtered HSQCNOESY spectrum of 0.2 mM active $^{13}C$-labeled $\alpha_M$I-domain with 1 mM unlabeled PTN-CTD. There are no observed side chain contacts between the proteins.

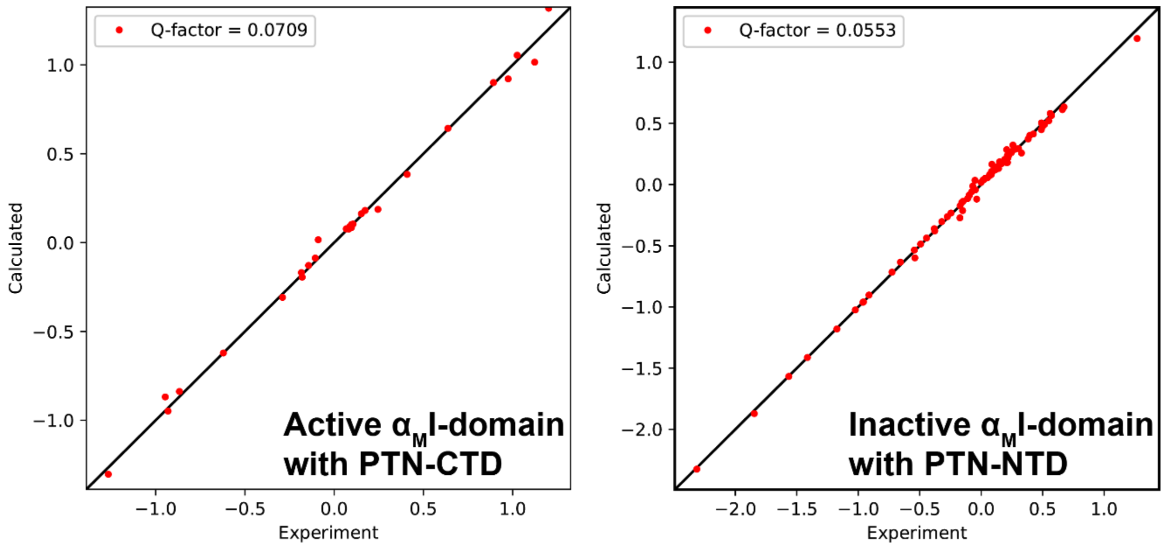


**Figure S9**. PCS fitting for active $\alpha_M$I-domain with PTN-CTD (left) and inactive $\alpha_M$I-domain with PTN-NTD (right). PDB 1IDO was used to fit the active $\alpha_M$I-domain PCS and PDB 1JLM was used to fit the inactive $\alpha_M$I-domain PCS. Presence of PTN domains did not change $\alpha_M$I-domain's PCS. This indicates the structures of inactive and active $\alpha_M$I-domain were not changed by PTN binding.

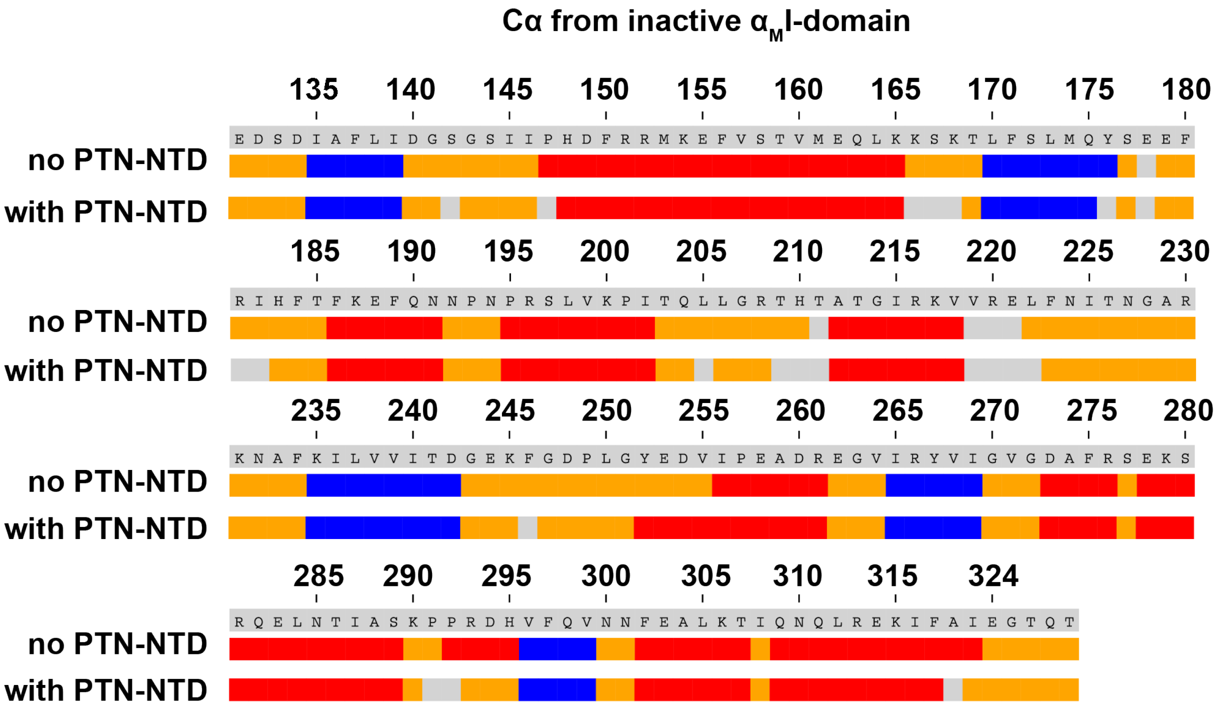


**Figure S10**. The predicted secondary structure of inactive $\alpha_M$I-domain calculated from Cα chemical shifts of the protein in the presence and absence of PTN-NTD. α helices are represented by red, β sheets are represented by blue, and coils are represented by orange. Residues with no assignment are shown as grey. The results show that the secondary structure of inactive $\alpha_M$I-domain remains the same when complexed with PTN-NTD.

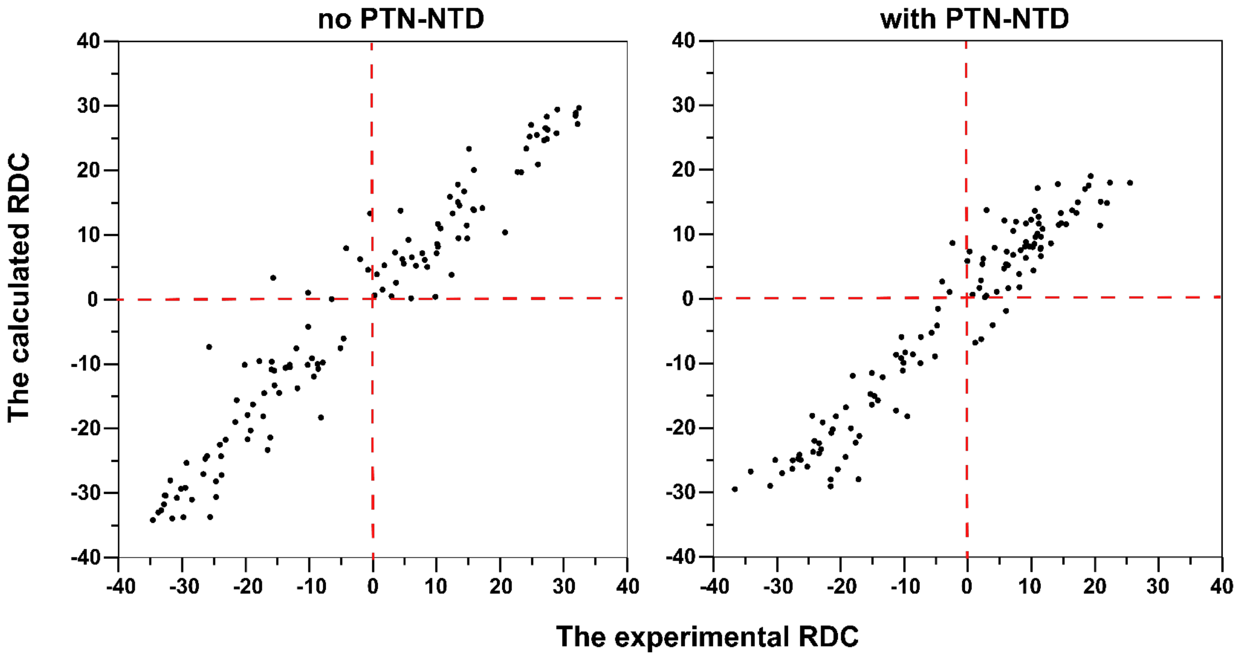


**Figure S11**. The correlation between the experimental and calculated RDC of inactive $\alpha_M$I-domain in the absence and presence of PTN-NTD. The results show that the experimental RDC of inactive $\alpha_M$I domain are well correlated with the crystal structure of inactive $\alpha_M$I-domain (PDB 1JLM) with a Q factor of 0.251. For inactive in the presence of PTN-NTD, the Q factor is 0.271. These data suggest the structure of $\alpha_M$I-domain did not undergo significant changes in the presence of ligands.

**Table S1**. The Kds of interaction between active $\alpha_M$I-domain and PTN domains as well as Glutamate.

| Signal | PTN | PTN-CTD | PTN-NTD | Glutamate |
|---|---|---|---|---|
| K245 | 0.07 ± 0.070 | 0.083 ±0.020 | 0.380 ± 0.202 | 7.004 ± 0.263 |
| F275 | 0.051 ± 0.026 | 0.074 ± 0.012 | 0.650 ± 0.253 | 6.036 ± 0.302 |
| G272 | 0.044 ± 0.045 | 0.168 ± 0.027 | 0.298 ± 0.081 | 6.459 ± 1.041 |
| G251 | 0.012 ± 0.027 | 0.085 ± 0.017 | 0.845 ± 0.278 | 5.367 ± 0.384 |
| S277 | 0.086 ± 0.034 | 0.091 ± 0.028 | 0.971 ± 0.486 | 5.315 ± 0.352 |
| G321 | 0.013 ± 0.014 | 0.085 ± 0.027 | 1.290 ± 0.528 | 4.798 ± 0.510 |
| R276 | 0.003 ± 0.031 | 0.062 ± 0.026 | 0.311 ± 0.026 | 7.370 ± 0.626 |
| TrendNMR | 0.350 ± 0.039 | 0.226 ± 0.020 | 6.250 ± 8.072 | 7.609 ± 0.587 |

**Table S2**. The Kd of the interaction between active $\alpha_M$I-domain and PTN-CTD mutants.

| Signal | PTN-CTD | E76QD78N | PTN-CTD-Δ tail | H95S | E98Q | E66QE98Q |
|---|---|---|---|---|---|---|
| K245 | 0.083 ± 0.020 | 0.147 ± 0.031 | 0.012 ± 0.049 | 0.074 ± 0.030 | 0.787 ± 1.381 | 1.010 ± 0.103 |
| F275 | 0.074 ± 0.020 | 0.164 ± 0.038 | 0.189 ± 0.025 | 0.088 ± 0.040 | 1.192 ± 0.404 | 0.758 ± 0.345 |
| G272 | 0.168 ± 0.027 | 0.235 ± 0.023 | 0.233 ± 0.039 | 0.075 ± 0.024 | 1.2515 ± 0.280 | 3.245 ± 2.860 |
| G251 | 0.085 ± 0.017 | 0.178 ± 0.026 | 0.171 ± 0.059 | 0.256 ± 0.155 | 1.161 ± 0.211 | 1.636 ± 0.827 |
| S277 | 0.091 ± 0.028 | 0.246 ± 0.079 | 0.218 ± 0.029 | 0.073 ± 0.015 | 0.979 ± 0.127 | 0.987 ± 0.235 |
| G321 | 0.085 ± 0.027 | 0.190 ± 0.058 | 0.677 ± 0.241 | 0.427 ± 0.088 | 0.302 ± 0.042 | 0.302 ± 0.042 |
| R276 | 0.062 ± 0.026 | 0.082 ± 0.018 | 0.126 ± 0.015 | 0.042 ± 0.013 | 0.286 ± 0.026 | 0.172 ± 0.030 |
| TrendNMR | 0.226 ± 0.019 | 0.237 ± 0.030 | 0.226 ± 0.019 | 0.319 ± 0.112 | 2.019 ± 0.837 | 5.230 ± 6.330 |

**Table S3**. Active $\alpha_M$I-domain backbone amide chemical shift changes produced by PTN-CTD mutations. Residues that are part of MIDAS are highlighted in yellow.

| | E76QD78N | PTN-CTD-Δ tail | H95S | E98Q | E66QE98Q |
|---|---|---|---|---|---|
| 132 | 0.011 | 0.010 | 0.018 | 0.016 | 0.004 |
| 140 | 0.014 | 0.002 | 0.010 | 0.014 | 0.014 |
| 141 | 0.006 | 0.005 | 0.011 | 0.007 | 0.006 |
| 142 | 0.008 | 0.034 | 0.071 | 0.045 | 0.043 |
| 143 | 0.047 | 0.033 | 0.065 | 0.066 | 0.071 |
| 144 | 0.000 | 0.033 | 0.222 | 0.138 | 0.159 |
| 145 | 0.006 | 0.006 | 0.054 | 0.051 | 0.053 |
| 146 | 0.022 | 0.020 | 0.083 | 0.033 | 0.073 |
| 149 | 0.006 | 0.004 | 0.008 | 0.018 | 0.016 |
| 150 | 0.005 | 0.000 | 0.003 | 0.006 | 0.012 |

| | | | | | |
|---|---|---|---|---|---|
| 151 | 0.007 | 0.008 | 0.015 | 0.012 | 0.015 |
| 152 | 0.040 | 0.022 | 0.048 | 0.044 | 0.043 |
| 153 | 0.002 | 0.012 | 0.015 | 0.012 | 0.009 |
| 154 | 0.006 | 0.004 | 0.003 | 0.008 | 0.004 |
| 155 | 0.010 | 0.012 | 0.016 | 0.016 | 0.020 |
| 156 | 0.019 | 0.010 | 0.047 | 0.052 | 0.047 |
| 160 | 0.015 | 0.007 | 0.022 | 0.009 | 0.013 |
| 165 | 0.009 | 0.005 | 0.147 | 0.011 | 0.009 |
| 170 | 0.004 | 0.007 | 0.023 | 0.012 | 0.026 |
| 171 | 0.035 | 0.013 | 0.007 | 0.018 | 0.029 |
| 172 | 0.014 | 0.014 | 0.029 | 0.008 | 0.007 |
| 182 | 0.002 | 0.004 | 0.007 | 0.006 | 0.009 |
| 188 | 0.006 | 0.009 | 0.007 | 0.004 | 0.006 |
| 189 | 0.006 | 0.006 | 0.005 | 0.007 | 0.000 |
| 190 | 0.012 | 0.014 | 0.012 | 0.016 | 0.007 |
| 194 | 0.008 | 0.006 | 0.012 | 0.012 | 0.011 |
| 198 | 0.013 | 0.009 | 0.007 | 0.017 | 0.012 |
| 203 | 0.018 | 0.024 | 0.015 | 0.147 | 0.027 |
| 204 | 0.006 | 0.007 | 0.003 | 0.048 | 0.002 |
| 205 | 0.016 | 0.018 | 0.011 | 0.015 | 0.013 |
| 206 | 0.039 | 0.053 | 0.023 | 0.075 | 0.044 |
| 207 | 0.022 | 0.006 | 0.060 | 0.053 | 0.144 |
| 213 | 0.007 | 0.002 | 0.004 | 0.011 | 0.013 |
| 215 | 0.003 | 0.006 | 0.002 | 0.016 | 0.009 |
| 224 | 0.003 | 0.002 | 0.006 | 0.011 | 0.010 |
| 226 | 0.001 | 0.006 | 0.006 | 0.012 | 0.006 |
| 227 | 0.002 | 0.002 | 0.002 | 0.020 | 0.014 |
| 228 | 0.006 | 0.003 | 0.005 | 0.012 | 0.011 |
| 229 | 0.001 | 0.004 | 0.001 | 0.013 | 0.019 |
| 230 | 0.002 | 0.005 | 0.001 | 0.010 | 0.002 |
| 231 | 0.002 | 0.003 | 0.001 | 0.007 | 0.003 |
| 233 | 0.001 | 0.001 | 0.005 | 0.006 | 0.007 |
| 242 | 0.006 | 0.004 | 0.020 | 0.023 | 0.020 |
| 243 | 0.011 | 0.012 | 0.035 | 0.035 | 0.035 |
| 244 | 0.004 | 0.002 | 0.052 | 0.010 | 0.006 |
| 245 | 0.000 | 0.004 | 0.049 | 0.037 | 0.050 |
| 246 | 0.000 | 0.000 | 0.000 | 0.048 | 0.042 |
| 247 | 0.010 | 0.008 | 0.000 | 0.000 | 0.000 |
| 250 | 0.004 | 0.012 | 0.010 | 0.075 | 0.007 |
| 251 | 0.003 | 0.003 | 0.008 | 0.005 | 0.010 |
| 252 | 0.000 | 0.003 | 0.004 | 0.006 | 0.000 |
| 253 | 0.001 | 0.003 | 0.014 | 0.004 | 0.008 |
| 254 | 0.000 | 0.033 | 0.036 | 0.039 | 0.005 |
| 258 | 0.005 | 0.002 | 0.005 | 0.008 | 0.008 |
| 259 | 0.001 | 0.005 | 0.008 | 0.007 | 0.008 |
| 260 | 0.002 | 0.003 | 0.004 | 0.007 | 0.005 |

| | | | | | |
|---|---|---|---|---|---|
| 261 | 0.002 | 0.003 | 0.004 | 0.011 | 0.009 |
| 262 | 0.005 | 0.008 | 0.008 | 0.043 | 0.010 |
| 263 | 0.003 | 0.003 | 0.004 | 0.011 | 0.008 |
| 264 | 0.012 | 0.005 | 0.015 | 0.011 | 0.004 |
| 266 | 0.025 | 0.007 | 0.022 | 0.019 | 0.020 |
| 267 | 0.009 | 0.006 | 0.009 | 0.005 | 0.005 |
| 272 | 0.006 | 0.005 | 0.018 | 0.041 | 0.026 |
| 273 | 0.001 | 0.004 | 0.015 | 0.022 | 0.020 |
| 274 | 0.003 | 0.003 | 0.005 | 0.003 | 0.003 |
| 275 | 0.001 | 0.002 | 0.015 | 0.024 | 0.065 |
| 276 | 0.009 | 0.005 | 0.007 | 0.028 | 0.009 |
| 277 | 0.001 | 0.002 | 0.015 | 0.009 | 0.010 |
| 278 | 0.026 | 0.009 | 0.006 | 0.013 | 0.028 |
| 279 | 0.001 | 0.002 | 0.014 | 0.010 | 0.011 |
| 280 | 0.002 | 0.001 | 0.030 | 0.009 | 0.006 |
| 281 | 0.006 | 0.008 | 0.015 | 0.033 | 0.027 |
| 283 | 0.006 | 0.000 | 0.008 | 0.018 | 0.017 |
| 287 | 0.003 | 0.006 | 0.003 | 0.007 | 0.003 |
| 288 | 0.008 | 0.006 | 0.004 | 0.002 | 0.006 |
| 289 | 0.004 | 0.012 | 0.008 | 0.002 | 0.020 |
| 296 | 0.007 | 0.006 | 0.018 | 0.033 | 0.053 |
| 297 | 0.004 | 0.004 | 0.001 | 0.011 | 0.004 |
| 298 | 0.006 | 0.005 | 0.005 | 0.008 | 0.007 |
| 311 | 0.000 | 0.006 | 0.005 | 0.011 | 0.008 |
| 312 | 0.014 | 0.017 | 0.010 | 0.013 | 0.011 |
| 313 | 0.003 | 0.008 | 0.006 | 0.009 | 0.009 |
| 314 | 0.005 | 0.002 | 0.008 | 0.017 | 0.011 |
| 316 | 0.007 | 0.014 | 0.014 | 0.007 | 0.008 |
| 317 | 0.005 | 0.007 | 0.018 | 0.038 | 0.027 |
| 318 | 0.006 | 0.008 | 0.017 | 0.017 | 0.013 |
| 319 | 0.002 | 0.001 | 0.005 | 0.009 | 0.007 |
| 320 | 0.002 | 0.004 | 0.012 | 0.012 | 0.010 |
| 321 | 0.000 | 0.000 | 0.008 | 0.010 | 0.010 |
| 322 | 0.002 | 0.003 | 0.004 | 0.013 | 0.006 |
| 323 | 0.002 | 0.002 | 0.005 | 0.003 | 0.007 |
| 324 | 0.005 | 0.008 | 0.005 | 0.008 | 0.006 |

**Table S4**. Intermolecular distance constraints used in HADDOCK docking of PTN-NTD onto inactive $\alpha_M$I-domain.

| PTN-NTD Atom | | $\alpha_M$I-domain Atom | | Distance (Å) |
|---|---|---|---|---|
| Residue Num. | Atom Name | Residue Num. | Atom Name | |
| G31 | HA# | R261 | HA | 2.7±0.5 |
| L32 | HD# | I265 | HD1# | 2.7±0.5 |
| L32 | HD# | I265 | HG2# | 2.7±0.5 |
| L32 | HD# | G263 | HA# | 2.7±0.5 |
| L32 | HD# | D260 | HA | 2.7±0.5 |
| T34 | HG2# | P291 | HD# | 2.7±0.5 |
| T34 | HG2# | K290 | HE# | 2.7±0.5 |
| T50 | HG2# | P291 | HD# | 2.7±0.5 |
| T26 | HG2# | K290 | HE# | 2.7±0.5 |
| R52 | HD# | I265 | HD1# | 2.7±0.5 |

**Table S5**. Intermolecular distance constraints used in HADDOCK docking of PTN-CTD onto active $\alpha_M$I-domain.

| Metal | PTN-CTD Atom | | $\alpha_M$I-domain Atom | | Distance (Å) |
|---|---|---|---|---|---|
| | Residue Num. | Atom Name | Residue Num. | Atom Name | |
| $Mg^{2+}$ | | | 209 | OG1 | 2.12 |
| $Mg^{2+}$ | | | S144 | OG | 2.58 |
| $Mg^{2+}$ | | | S142 | OG | 2.03 |
| $Mg^{2+}$ | E98 | OE# | | | 2.20 |
| | E98 | HG# | S144 | H | 3.0±0.5 |
| | E98 | HG# | G143 | H | 3.0±0.5 |
| | E98 | HG# | I145 | H | 3.0±0.5 |
| | E98 | HG# | R208 | H | 3.0±0.5 |
| | A93 | HB# | S144 | H | 3.0±0.5 |
| | A93 | HB# | R208 | H | 3.0±0.5 |
